# On the mechanical regulation of epithelial tissue homeostasis

**DOI:** 10.1101/2021.04.10.439119

**Authors:** Sara Kaliman, Maxime Hubert, Carina Wollnik, Lovro Nuić, Damir Vurnek, Simone Gehrer, Jakov Lovrić, Diana Dudziak, Florian Rehfeldt, Ana-Sunčana Smith

## Abstract

Despite recent efforts to understand homeostasis in epithelial tissues, there are many unknowns surrounding this steady state. It is considered to be regulated by mechanoresponse, but unlike for single cells, this remains heavily debated for tissues. Here, we show that changes in matrix stiffness induce a non-equilibrium transition from tubular to squamous Madin-Darby Canine Kidney II tissues. Nonetheless, despite different cell morphologies and densities, all homeostatic tissues display equivalent topologies, which, hence, must be actively targeted and regulated. On the contrary, the mechanoresponse induces dramatic changes in the large-scale organization of the colonies. On stiff gels, this yields an unreported cooperative state of motile cells displaying higher densities than in the arrested homeostatic state. This suggests a more complex relation between cell density and motility than previously anticipated. Our results unequivocally relate the mechanosensitive properties of individual cells to the evolving macroscopic structures, an effect that could be important for understanding the emergent pathologies of living tissues.

## I. Introduction

Homeostasis [1] denotes maintenance of the morphology and metabolic functions in differentiated tissues. It is preceded by tissue growth or regeneration in which complex tissue architecture emerges on multiple length scales [2]. It spans from the size of a cell, to the mesoscopic level with structures such as crypts [3], to the macroscopic formation of entire organs. In epithelium, major strategies of homeostatic control are contact inhibition of proliferation [4] and locomotion [5], as well as cell extrusions [6], which are all dependent on the actual cell density. As such, the homeostatic steady state H_PL_ (inhibition of Proliferation and Locomotion), with its characteristic structure, is cooperative by nature [7], and a result of non-equilibrium self-organization, a process that has not been fully understood so far. In the biomedical context, however, the homeostatic states, with their characteristic cell densities and shapes, have been very well characterized. Consequently, deviations from typical phenotypes, both on the cellular and compartment level, are often considered important in the diagnosis of a broad range of diseases [8]. Nonetheless, the emergent tissue properties, as well as the biochemical and biophysical regulations of the homeostatic steady state are still subject to intense research [9–12].

Besides the complex biochemical signaling sequences involved in homeostasis [13], the H_PL_ architecture is tightly regulated by physical forces that are generated within and between the cells, as well as by the interactions with the extracellular matrix, as part of mechanoresponse [14, 15]. In individual adherent cells, the mechanosensitive force balance is strongly coupled with the cell shape, motility, nuclear positioning, and many other processes [10, 16–21]. During development, mechanoresponse is crucial for the compartmentalization of tissues and the development of organs. For example, the control of human epiblast and amnion development was found to be partly determined by the mechanical properties of the niche-like environment [22, 23]. In homeostatic tissues, however, the role of mechanosensing is still unclear. To the contrary, there is growing evidences that a number of pathological conditions involve changes in stiffness [17, 24, 25] and viscosity [26, 27] of the extracellular matrix. In cancer, for example, matrix stiffening has been related to the epithelial-mesenchymal transition and the significant increase of cell motility yielding metastasis [28]. However, mechanistic studies in model systems, involving the modulation of matrix stiffness, provided contradictory evidences regarding the properties and the topology of the H_PL_ steady state, the shapes of constitutive cells, as well as the macroscopic self-compartmentalization in growing colonies [29–36].

In order to resolve this debate and provide a deeper insight regarding the influence of cell mechanosensitivity at the microscopic and macroscopic levels, we perform here a systematic study of the effects of mechanosensing on the growth, self-assembly and homeostasis of a model epithelium. We take advantage of fully controlled, biomimetic environment to clearly delineate the mechanoresponse on different length and time scales, clearly demonstrating that both the shapes of cells and the compartmentalization of tissue during development respond to changes in the mechanical properties of the environment. Interestingly however, we find that the topology of the homeostatic tissue is independent of the stiffness, despite variations in density.

## II. Mechanoresponse of Cells Within the Homeostatic Tissue

We grow Madin-Darby Canine Kidney II (MDCK-II) tissues on substrates with systematically varying stiffness. The substrates are glass or polyacrylamide (PA) gels with a Young elastic modulus *E* = 0.6, 3, 5, 11, and 21 kPa (*δE* = ±0.3 kPa, see Appendix A for methodological details) covalently and uniformly coated with collagen-I. This change in substrate stiffness should result in different stress-generation patterns, hence altering the tissue steady state. To sample sufficient statistics on different length scales, we produce centimeter-wide monolayers, starting from a droplet containing approximately 30,000 cells. All colonies are grown for 4-6 days in controlled conditions, upon which the homeostatic state H_PL_ is fully formed in the central self-assembled compartment of the tissue (see Fig. 6 and Fig. 7). In H_PL_, if any proliferation occurs, it is balanced by the apoptosis rate, such that the mean density no longer depends on time. Furthermore, no net locomotion is observed (see Fig. 6).

**FIG. 1.**
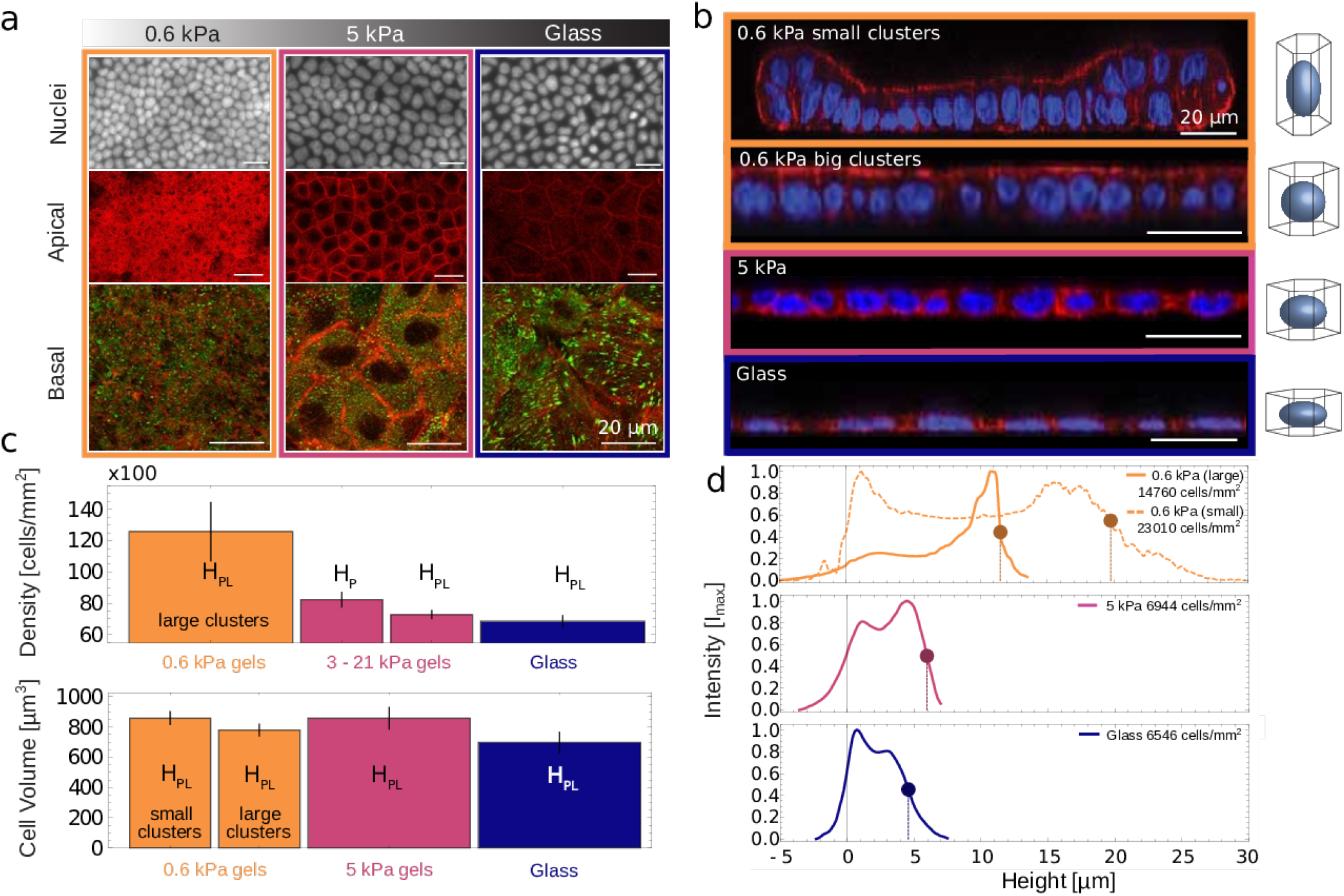
Characteristics of the homeostatic state of the tissue as a function of the substrate stiffness. Tissues are grown on soft gels (orange colored frames and lines), hard gels (pink) as well as on glass (blue) (a) Characteristic confocal images in the plan of the monolayers fixed early in the 4th day after seeding. The top panel presents Hoechst stained cell nuclei. In the middle apical phalloidin stained actin is imaged. The bottom panel shows paxilin stained using secondary antibodies in green and actin in red to visualize focal adhesions. (b) Reconstructed slice through the monolayers showing columnar MDCK-II tissue on soft gels, cuboidal tissue on hard gels and squamous tissue on glass substrates. Actin is shown in red and nuclei in blue. In small colonies on soft gels are characterized by a monolayer area smaller than 2.8 × 10^−2^ mm^2^, and finite size effects still play a role. Larger colonies are imaged away from the edge. (c) Bar charts of the average cell density (top) and volume (bottom). Error bars indicate standard deviations from measurements in different colonies grown under identical conditions. This deviation is however similar to standard deviations in density within a single colony (see Appendix B for details). Unlike cell volumes in different tissues, cell density shows statistically significant differences even between hard gels and glass. However, the error in the measurement of the volume, obtained from the propagation of uncertainties in measuring density and height, is of the order of 10%. Density here was calculated from approximately 500 × 10^3^ cells at 0.6 kPa, from 208 × 10^3^ cells grown in tissues on substrates with stiffness between 3 and 21 kPa and from 27 × 10^3^ cells on glass (see Appendix B2). Height was calculated for 100 (glass) to 350 cells (soft gels) depending on the density considered. The notations H_PL_ and H_P_ stands for the homeostatic state with inhibition of proliferation and locomotion and for the homeostatic state with inhibition of proliferation respectively. (d) Average intensity of the stained actin as a function of the distance from the substrate. The height of the monolayer is determined as the difference in heights between the two inflection points in the intensity curves. For simplicity, all inflection points at the basal surface are co-aligned. The circles indicate the apical surface. The density in the presented segment where the volume is measured is indicated in each graph.

**FIG. 2.**
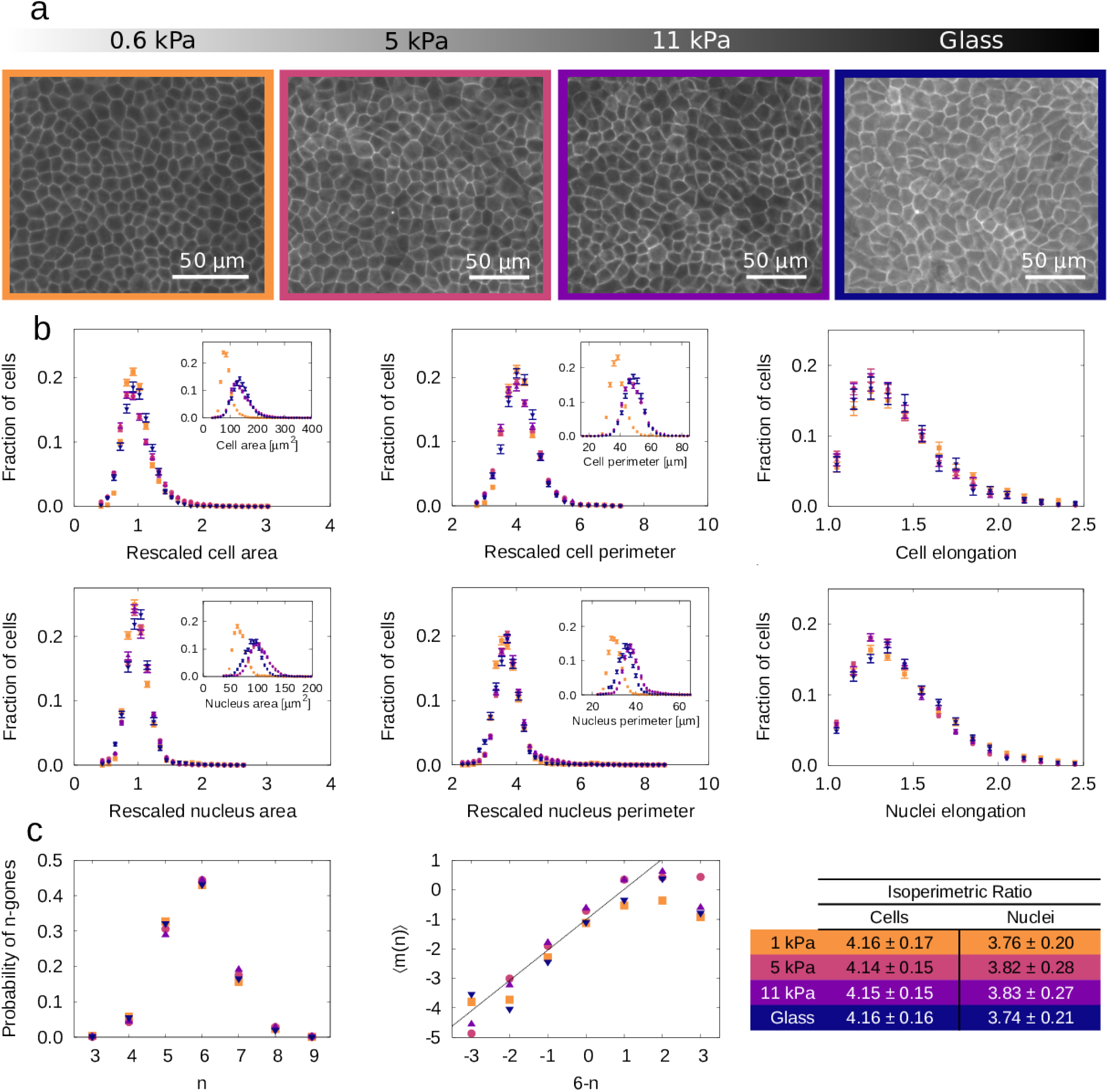
Structure and topology of tissues in the homeostatic state. (a) Images of *β*-catenin stained cell membranes in the steady state H_PL_ (inhibited for proliferation and locomotion). The magnification in different frames is chosen such that cells appear equally large, to visualize the morphological re-scaling. (b) Distributions of morphological measures of cells obtained through set Voronoi tessellations (top row) and nuclei (bottom row). For soft gels (orange) 4257 cells are extracted from large colonies to avoid finite size effects. For tissues grown on 5 kPa (pink), 11 kPa (purple), and glass (blue) substrates 9095, 5244 cells and 2575 cells are considered. The distributions of re-scaled measures (each length divided by the square root of the mean area) are provided as main graphs, while the insets show the original data. Distributions for area (left), perimeters (middle) and elongations (right) are shown. Elongations are extracted by calculating the principle axis of inertia of the relevant object (cell or nucleus), and finding their ratio. Error bars correspond to the standard deviation on 50 independent subsets of cells (see Appendix C). (c) Conservation of topological measures relative to tissues grown on different substrates. The color code is same as throughout the figure. Distribution of cells with *n* sides (i.e. neighbors) is on the left, while the Aboav-Weaire’s law here measure through the parameter 〈*m*(*n*)〉 = −(6 − *μ*(*n*)*n* − *σ*(*n*)^2^. The isoperimetric ratio for the average cell and nuclei shapes are provided in the table on the right.

**FIG. 3.**
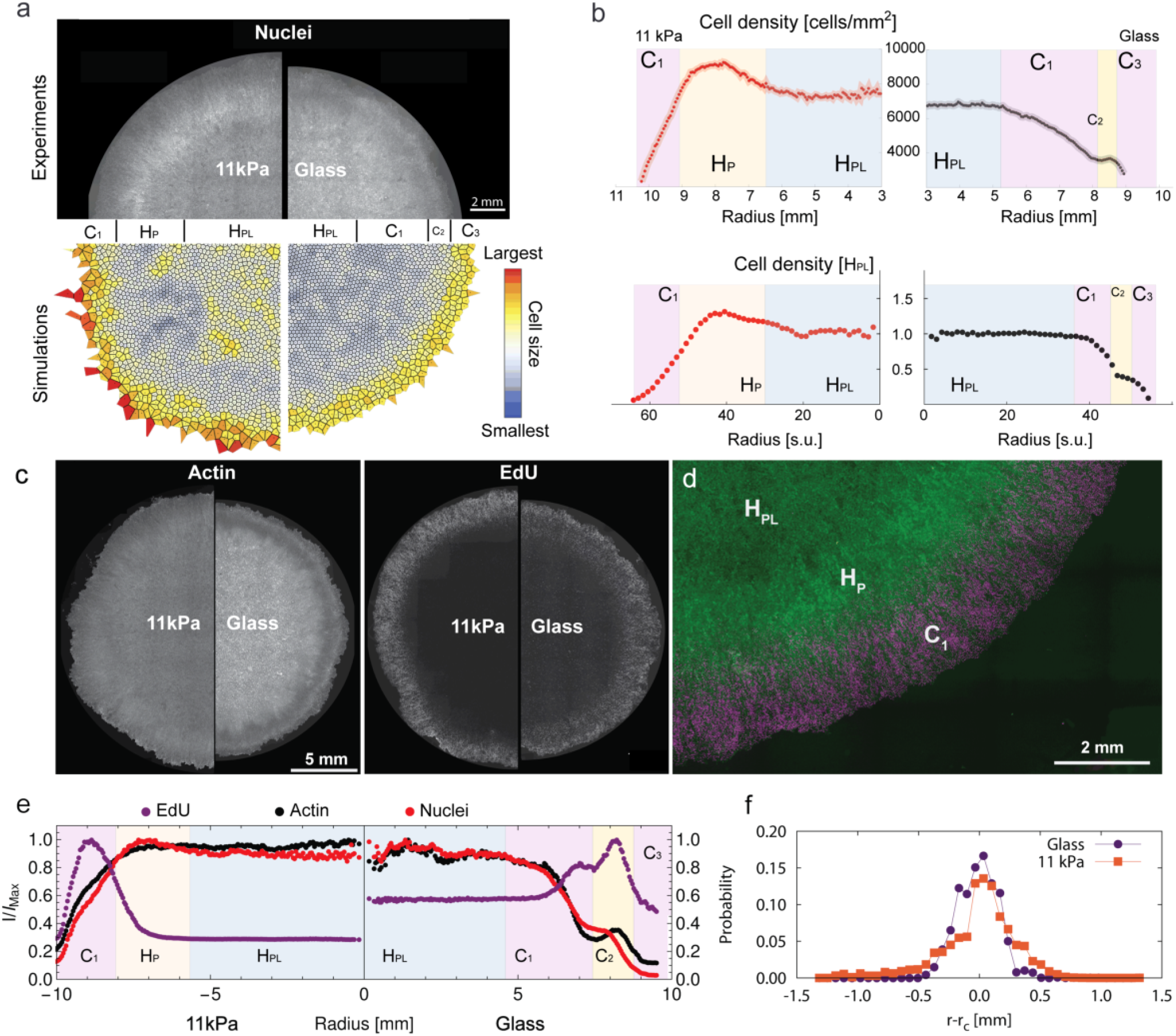
Macroscopic organization of the tissue surrounding the homeostatic state on hard gels and glass. H_PL_ and H_P_ stands for the homeostatic state with inhibition of proliferation and locomotion and for the homeostatic state with inhibition of proliferation respectively. Additional compartments are denoted with *C*_*n*_, as defined in the main text. (a) Hoechst stained cell nuclei throughout the tissues grown for 6 days (top panel) are compared to simulations (bottom panel). Colonies on hard gels (left panel) are compared to ones developed on glass substrates (right panel). Scale bar indicates 2 mm. Various compartments are indicated in between the simulation and experimental diagrams. (b) Experimental density profiles obtained from the linear relation between density and light intensity (top row) on gels (red curve on the left) and glass (black curves on the right), more details in Appendix D1. Simulation counterparts are shown in the bottom row. (c) Experimental cell clusters on hard gels and glass substrates stained for actin and EdU, showing only residual proliferation in the H_PL_ state (d) Segment of a tissue grown on hard gels showing no proliferation within the H_PL_ and H_P_ compartments. The purple, EdU stained nuclei are overlayed with the Hoechst stain in green. (e) Intensity profiles for actin (black), nuclei (red) and EdU (purple) through the colonies grown on hard 11 kPa gels and glass. (f) Tissue roughness evaluated by the normalized distributions of the deviations of the edge from the average shape.

**FIG. 4.**
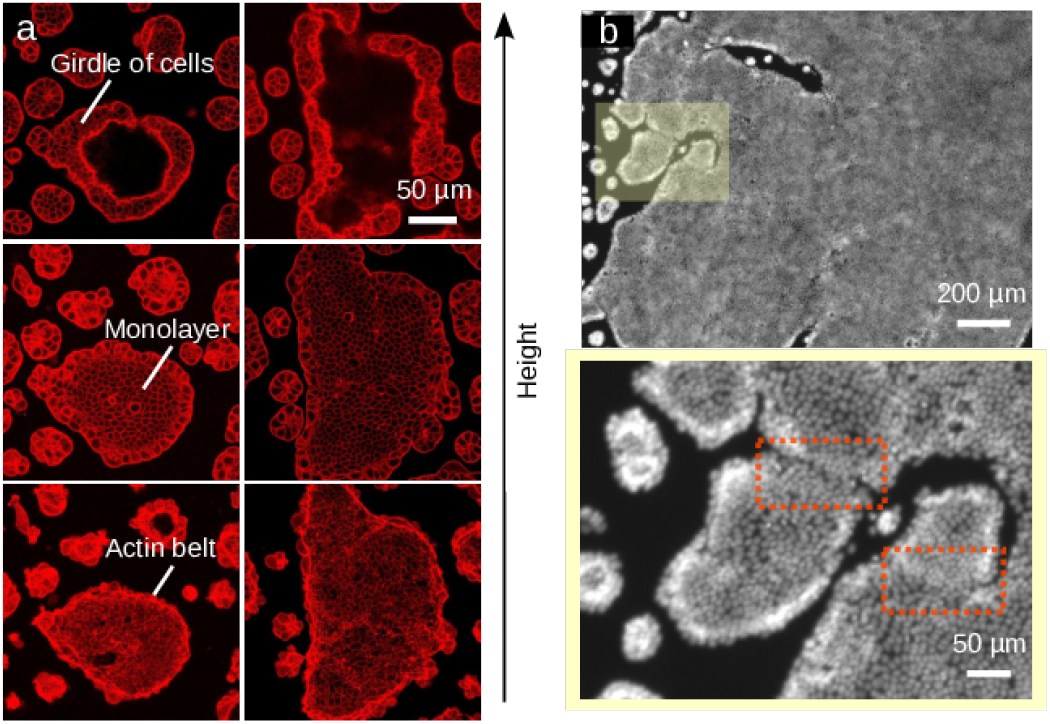
Tissues grown on soft gels. (a) Actin stained cells within clusters grown on soft substrates (E = 0.6 kPa). Three confocal slices of the same colony are provided focusing on the basal side with the actin belt (bottom), central monolayer (middle), and the multilayerd girdle of cells (top). (b) Epifluorescence image of nuclei stained with Hoechst dye of a tissue grown on soft 0.6 kPa gels. The yellow box in the top panel is enlarged in the bottom. The girdle and the monolayer can be clearly discerned. Spherical agglomerates, small colonies and a segment of the large colony is shown simultaneously. In the bottom panel line-like scars due to the merging of two colonies are boxed.

**FIG. 5.**
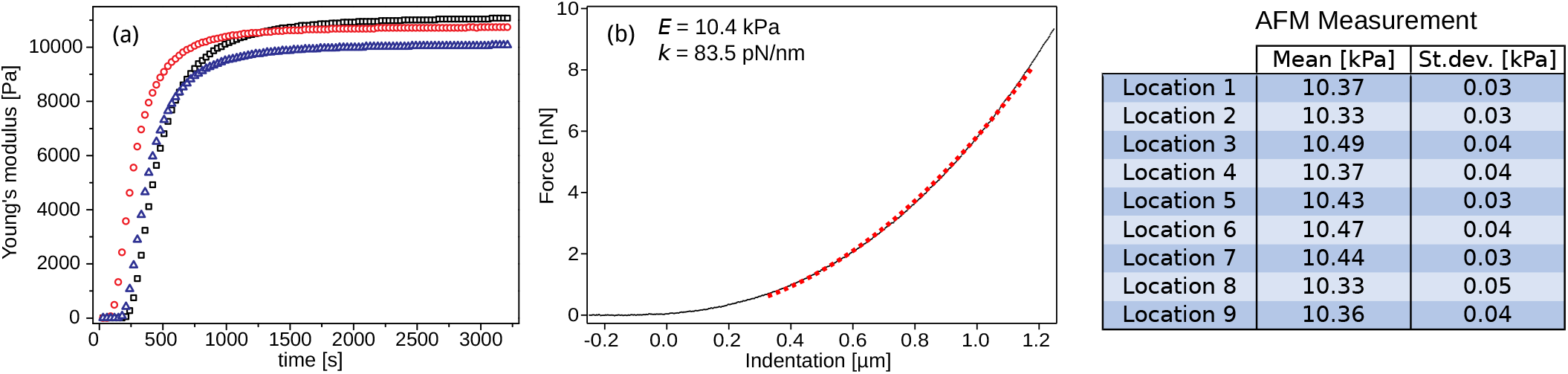
Characterization of the gels stiffness. (a) Bulk rheology measurement of the Young’s elastic modulus *E* over time. Plateau values are 11.1 kPa (black squares), 10.7 kPa (red circles) and 10.1 kPa (blue triangles). This leads an average value *E* = 10.6 ± 0.3 kPa. (b) AFM measurement of the force distance-curve (black dots) and the fitted Hertz model red dashed line giving the Young’s modulus *E* of the gels as 10.4 kPa. The table shows the mean values of the fitted Young’s modulus *E* of 10 measurements at each of the 9 locations on the PA gel.

**FIG. 6.**
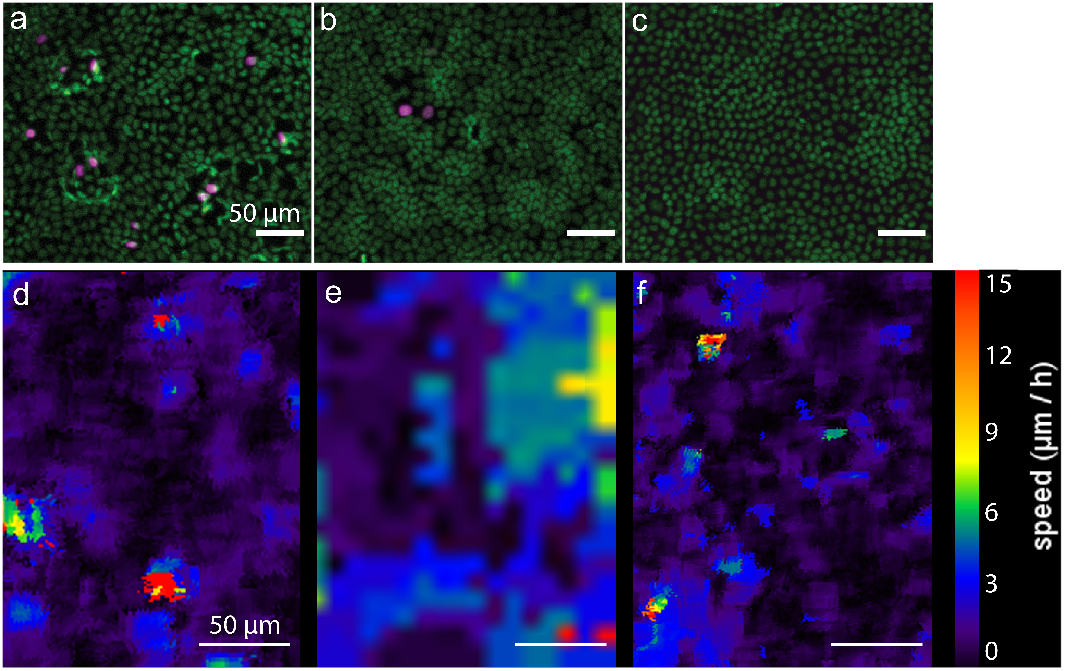
Contact inhibition with respect to proliferation and motion. Illustration of the contact inhibition for proliferation using EdU staining (top row) and for motion through PIV analysis (bottom row). (a,b,c) Images of EDU (purple) and nuclei (in green) stained H_PL_ for glass, hard gels and soft gels respectively, showing no cell divisions in the H_PL_. Both on glass and on gels, the majority of the division events were observed in a vicinity of a defect in a tissue. Scale-bar is 50 *μ*m. (d,e,f) PIV velocity fields within the H_PL_ showing low cell mobility both on glass, 20 kPa gels and soft gels respectively. (d) MDCK-II cells were seeded on a soft gel substrate with a Young modulus of 0.75 kPa in a 2 *μ*L dense droplet of 100 000 cells on a covalently bound collagen-I coated surface. Imaged 2.5 days after seeding. Pixel size 1.25 *μ*m, time step 15 min. (e) MDCK-II cells were seeded on a hard gel substrate with a Young modulus of 30 kPa in a 2 *μ*L dense droplet of 10 000 cells on a covalently bound collagen-I coated surface. Imaged 2.7 days after seeding. Pixel size 10 *μ*m, time step 20 min. (f) MDCK-II cells were seeded on a glass substrate with in a 2 *μ*L dense droplet of 10 000 cells on a covalently bound collagen-I coated surface. Imaged 10 days after seeding. Pixel size 0.886 *μ*m, time step 20 min.

**FIG. 7.**
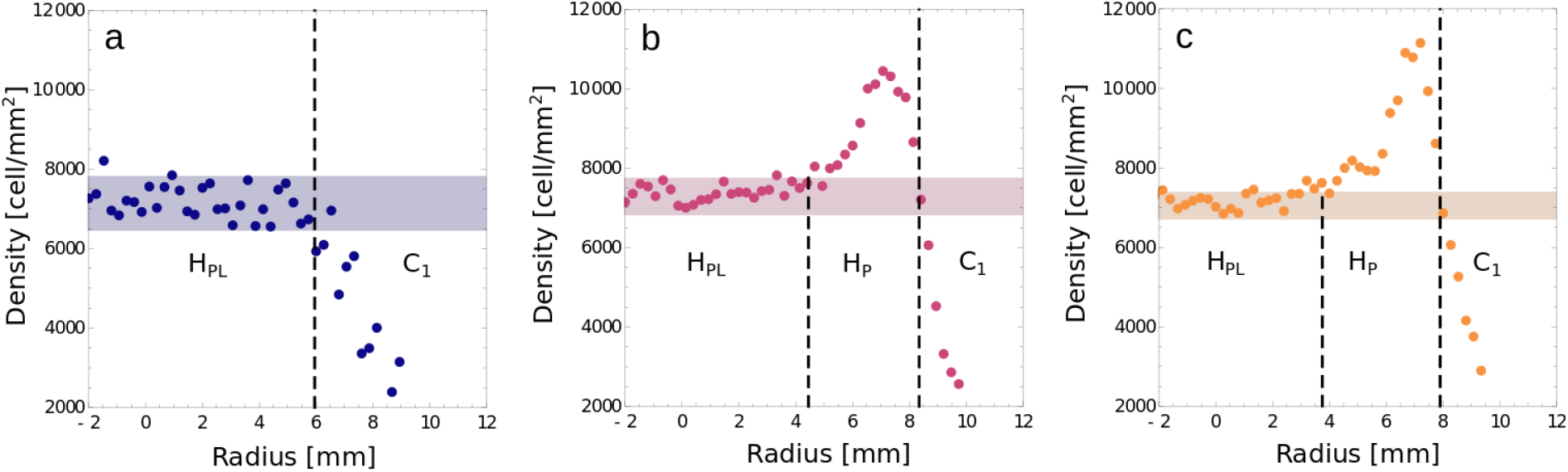
Spatial dependence of the density in the cluster. (a) Cluster on the glass substrate, (b) cluster on 11 kPa PA gels (c) cluster on 5 kPa gels, showing very similar features to the cluster grown on 11 kPa gels. These colonies belong to two different series of data in comparison to the ones shown in Fig. 1, but are mutually consistent. Horizontal lines represent two standard deviations around the mean cell density in the central region (2.3 mm radius). If the image segment is within those lines it is considered as bulk. If the density of a given image segment is below the bottom line the segment is considered as part of the edge region and segments with density higher than the upper line are considered as a part of the high density ring

Initially, we verify the mechanosensitivity of cells in homeostasis by imaging the subcellular cytoskeletal and adhesion structures as shown in Fig. 1a (for technical details and the biological function of stained proteins see Appendix A2). As reported in single cells, increasing the substrate rigidity yields larger focal adhesions and stronger actomyosin stress-fibers [21], but weaker apical tension (Fig. 1a) as evidenced by the modulation of apical and basal peaks in the actin vertical density profile (Fig. 1d).

On glass, the cell-substrate adhesion energy dominates with the actin cytoskeleton [14, 37] concentrating on the basal surface (Fig. 1d). The result is the appearance of squamous cells (height *h* = 4.6 ± 0.4 *μ*m, height-to-base length *h/r* = 0.35 ± 0.04) with a homeostatic density *ρ* (H_PL_) = 6860 ± 360 cells/mm^2^ (Fig. 1c). Under these conditions the basal tension is given by the formula from Hannezo et al [14] which reads 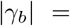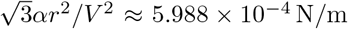 with *V* the cell volume and *α* ≈ 10^−24^ Jm^2^, the latter being the confinement energy of the cytoplasmic components.

On hard gels, the actin distribution displays both basal and apical maxima, indicating weakening of adhesion and strengthening of the apical tension (Fig. 1d). Consequently, the tissue becomes cuboidal (*h* = 5.9 ± 0.5 *μ*m, *h/r* = 0.47 ± 0.05). Surprisingly, the density of the homeostatic state *ρ* (H_PL_) = 7280 ± 260 cells/mm^2^ is slightly but consistently larger than on glass, yet statistically the same on gels of stiffness in the range of 3 kPa to 21 kPa (see Tab. 1). This trend is systematic over different series of experiments although the averages on glass and gels may differ of a few hundreds of cells per square micron, consistent with the differences noticed previously [38].

On soft gels, most actin is within the apical belt (Figs. 1b,d) to mediate a strong contractile tension Λ_*α*_, estimated by Hannezo et al [14] to be 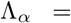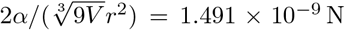. The emergent density is high *ρ* (H_PL_) = 12580 ± 1860 cells/mm^2^ and accompanied with a phase transition toward a columnar morphology (*h* = 11.5 ± 0.6 *μ*m, *h/r* = 1.20 ± 0.15). To accommodate, cell nuclei become highly asymmetric, with the major axis perpendicular to the substrate (Fig. 1b).

Furthermore, all homeostatic states have the same average cell volumes within the accuracy of the measurements (Fig. 1c). We can therefore conclude that cells in H_PL_ sense the stiffness of the underlying substrate by cooperatively adjusting their cell morphology as well as their nuclei shape and orientation. This is enabled by the modulations in adhesiveness and changes in the spatial distribution of actin.

## III. Universal Topology of Homeostatic States

We further investigate the organization of cells building the homeostatic states from a statistical perspective (technical details in Appendix C). Rather than extracting the morphological properties of cells directly from segmented images of the cell plasma membrane, which can be challenging, we approximate the cell shape using the Set-based Voronoi Tessellation, referred to as SVT in the reminder of this article. This method allows for generating very large and robust data sets as shown in Fig. 9. It relies on accurate segmentation of nuclei, which we achieve with our home-built post-treatment algorithm with an accuracy of 99%, even at the highest tissue densities [38]. As discussed in more detail in the Appendix C1, the accuracy of the SVT at the densities typically achieved for the H_P_ and H_PL_ states is high. The difference to direct segmentation of the membrane does not exceed 10% for any measure even when compared to a reference set extracted directly from high accuracy methods that do not support massive tissue imaging [38]. Using this approach we obtain distributions of cell and nuclei area, perimeter and elongation as shown in the insets of Fig. 2b.

**FIG. 8.**
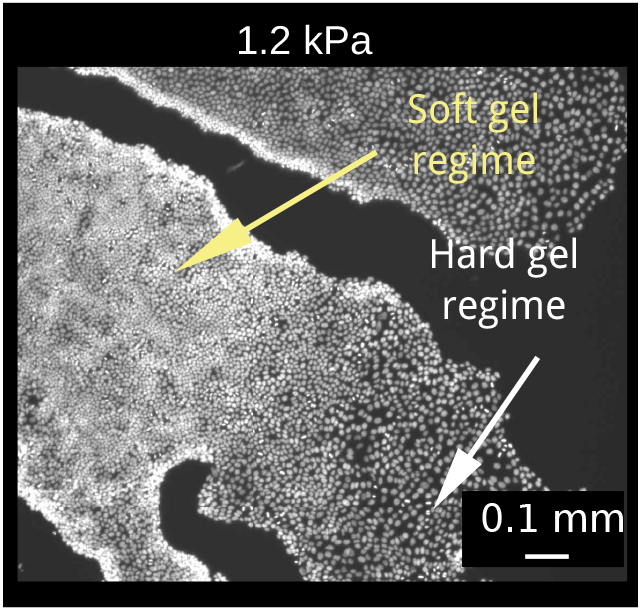
Tissues grown on soft gels with stiffness of 1.2 kPa. For slightly higher stiffness than 0.6 kPa, typically in the range of 1 kPa to 3 kPa, the tissue exhibits the behaviors expected on both soft gels and hard gels, as indicated on the picture by arrows. Image from [73].

**FIG. 9.**
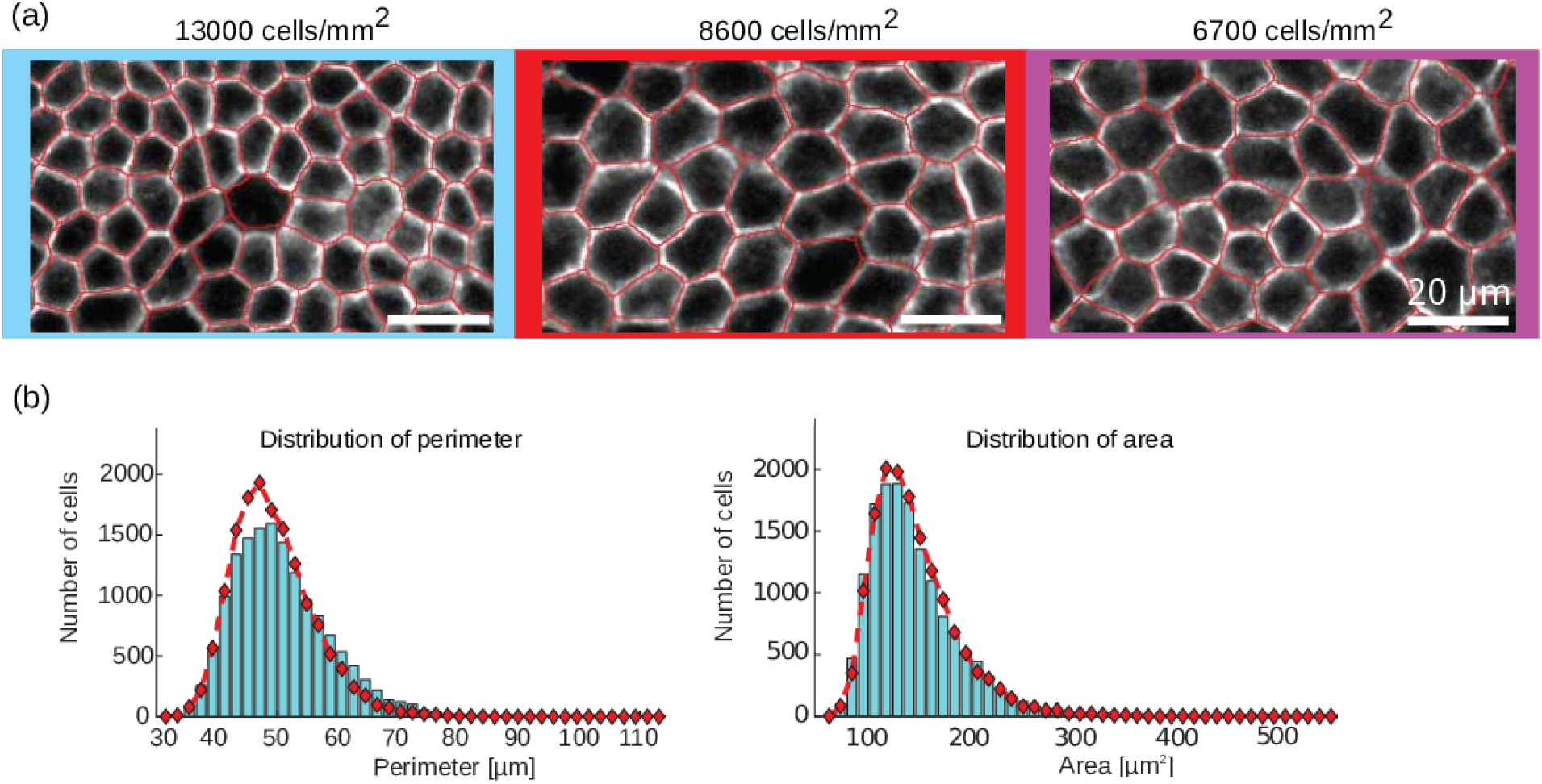
Morphological analysis of the tissue. (a) Comparison between pictures of the cell membranes stained with *β*-catenin and the set-based Voronoi tessellation (in red). (b) Distribution of cell perimeters and areas respectively measured by pictures of *β*-catenin stained membranes (bars) and by SVT (red diamonds) for cells grown on hard 11 kPa gels. These distributions comprise cells from all compartments of the tissue, including the edge compartments. Prior to the comparison, all segmented images of *β*-catenin and nuclei have been hand corrected for errors, as described in [38, 73].

For the purpose of establishing a unique metric between various tissues, all lengths are re-scaled by the square root of the average cell area in a particular tissue 〈*A*〉. This indeed leads to the collapse of data such that the distributions of area, perimeter and elongation for both cells and nuclei, show no appreciable differences as highlighted by the overlapping errorbars (Fig. 2b). Actually, the difference between two distributions of different substrates are comparable to differences between distributions generated from large subsets of cells obtained from the same substrate. This implies that the average cell, and the deviations from an average cells are, within the statistical accuracy of the data, self-similar among all tissues. Furthermore, all of the statistics driven from those distributions are within the accuracy of the measurements.The same conclusion holds for the cell nuclei, which is the distribution imposing the cell shapes in the plane of the monolayer.

Furthermore, to analyze the connectivity of the cells within the H_PL_ compartments of the tissues, we calculate the distributions of the number of neighbors *n* of all cells (Fig. 2c). Similarly to the case of morphological measures, we find that these distributions are statistically equal in all sets and insensitive to the substrate stiffness and therefore the cell density. Furthermore, these distributions are very similar to previously reported ones in different epithelium [39]. While the most common cell shape in all H_PL_ tissues is hexagonal (approximately 45%), the distributions of *n* are positively skewed. Actually the fractions of cells with five and seven neighbors are significant, with a larger pentagon component (30% pentagon compared to 20% heptagon). This asymmetry exists already in the random packings of ellipses at densities and elongations comparable to the nuclei density and shapes [40], albeit, in tissues, the disparity in the fractions of pentagons and heptagons is significantly larger.

Finally, to explore the topology of the tissues, we analyze the correlations between the number of neighbors of cell with *n* sides, and the average number of neighbors *m*(*n*) that cells adjacent to ones with *n* sides have. When this relation is linear, it is known as the Aboav-Weaire’s law [41–43] and states that

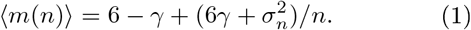

Here, 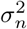 is the second central moment of the distribution of *n*, and *γ* is a constant that may decrease as 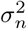 increases [44, 45], or may be independent of 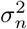 [46]. This relation has been commonly used to explore the topology of tessellations and arrangements in cellular assemblies [40, 47, 48]. In the case of homeostatic MDCK-II tissues, irrespective of the conditions in which they were grown, all tissues show the same linear dependence (Fig. 2c). This suggests that cells with fewer neighbors tend to have neighbors with more sides, and vice-versa. This trend is violated for cells with more than 8 or less than 4 neighbors. The linear relation shows also here a clear offset along the *y* axis. This suggest that the topology of the homeostatic MDCK-II, which is tissues emerges from the geometry of the nuclei spatial distribution, is actively maintained to achieve a particular connectivity.

The relation between geometry and topology can be further explored by calculating the correlations between the number of neighbors *n* and the average re-scaled area 〈*A*_*n*_〉 and re-scaled perimeter 〈*P*_*n*_〉 of a cell with *n* neighbors. These linear relations are known as the Lewis law [49–52] and Desh law [53], respectively. They suggest that large cells have a tendency to have more neighbors, while, inversely, small cells have a tendency to have less neighbors. Indeed, these linear relationships are also confirmed in MDCK-II tissues (see Fig. 10), suggesting again that geometrical elements, as emphasized by random packings [40] remain important even in tissues, even though the cell and nuclei shape distributions are strongly regulated in the homeostatic state.

**FIG. 10.**
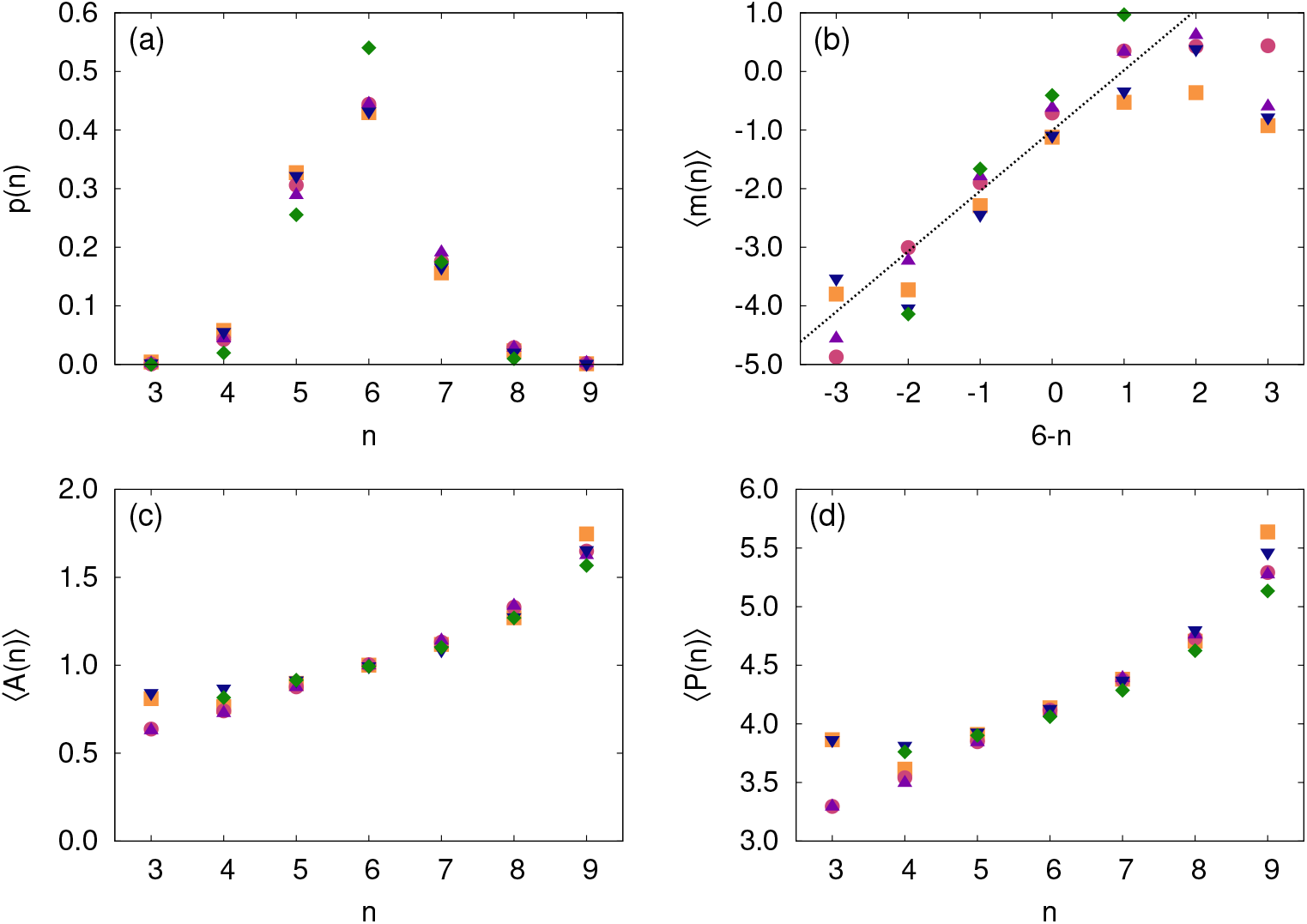
Topological equivalence of tissues on different substrates. (a) Histogram of the number of neighbors per cell for each substrate. (b) Aboav-Weaire’s law for each substrate. The black line seen on the figure is a fit on the linear part of the data. (c) Lewis’ law for each substrate. (d) Desh’s law for each substrate. The graphs consider tissues grown on all substrates: H_PL_ glass (blue down-triangle), H_PL_ 11 kPa gels (purple up-triangle), H_P_ 11 kPa gels (green diamonds), H_PL_ 5 kPa gels (pink circle) and H_PL_ 0.6 kPa gels (orange squares). For these two last laws, the area *A* of each cell and their perimeter *P* have been divided by the cell mean area and its square root respectively.

We thus conclude that tissue topology is the key property actively maintained in homeostasis leading to a well-defined spatial organization, by and large universal for all MDCK-II steady states. Universality of the topology is also reflected in the equivalence of the isoperimetric ratios of cells [54] and of cell nuclei, that remain independent of the matrix stiffness (table in Fig. 2c). This suggests that all homeostatic MDCK-II tissues are in mechanically equivalent steady states. The latter is, however, governed by very different contributions to the force balance, as reflected in different cell three-dimensional shapes and densities.

## IV. Macroscopic Effects of Mechanoresponse in Tissues Grown on Hard Gels and Glass

Finally, to bring additional insights about the role of mechanoresponse on the macroscopic scale (technical details in Appendix D), we investigate the structure of tissues supporting the homeostatic state. On glass and on hard gels, colonies are radially symmetric (Fig. 3a), and the H_PL_ state develops in the center (flat segment of the density profiles in top panels of Fig. 3b).

In the radial direction, after the H_PL_ state and towards to tissue outer rim, three more compartments are reported. We designate these compartments *C*_1_, *C*_2_ and *C*_3_. The large *C*_1_ compartment is characterized by cells which become motile, divide and their density continuously drops until the *C*_2_ compartment. The appearance of the latter is characterized by a change of curvature of the density profile (Fig. 3b,e). In *C*_2_ the cell density increases, while the basal actin shows stronger signal denoting stronger adhesion to the substrate compared to neighboring *C*_1_ and edge *C*_3_ compartment. Compartment *C*_3_ consists of a large number of cells that transiently adopt the phenotype of leader-cells [55] with extended lamellipodia (see Fig. 15). Cells in *C*_3_ commonly exchange their neighborhood, while the average speed of cells saturates (see Fig. 13).

**FIG. 11.**
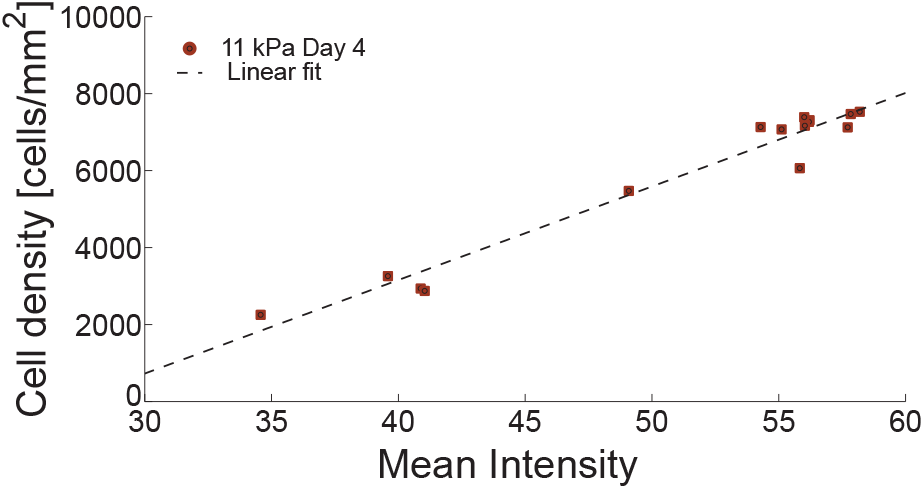
Relation between cell number density and mean intensity of the confocal images. During image post-treatment, cell clusters are divided in equidistant rings with the central region is a circle with a radius of 350 *μ*m. The image segments taken from the different regions in the cell clusters are analyzed and a linear mapping between average cell density and image intensity is found for each cluster. The figure illustrates the linear mapping extracted from the experiments realized on day 4 for tissues grown on 11 kPa gels.

**FIG. 12.**
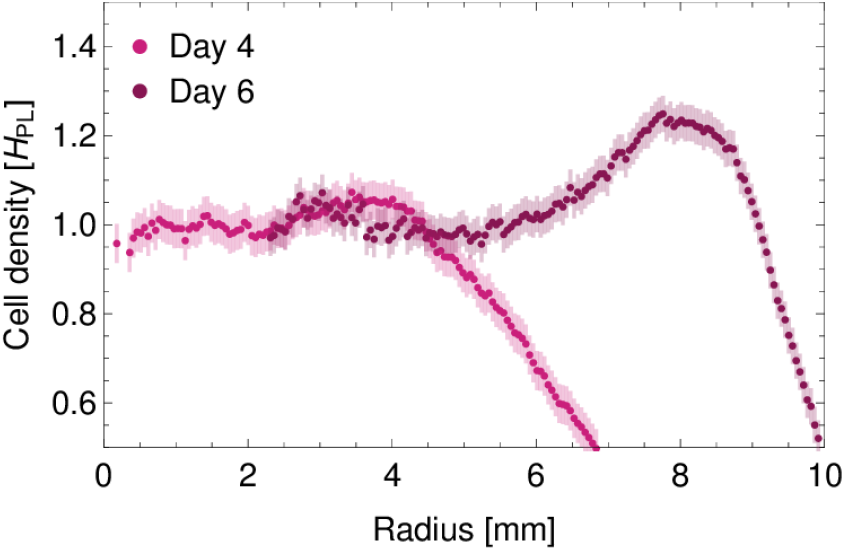
Mobility in the H_P_ compartment. Evolution of the density profile as a function of time for a tissue grown on hard gel, at day 4 (light pink) and at day 6 (dark pink). The weak local maximum at day 4 is seem to move from 4 mm to the center to 8 mm to the center at day 6, evidencing the motion of cell within the H_P_ state. Error bars corresponds to a typical variation of ±300 cells on the measurement of the density in the segment. The data in the first 2 mm from Day 6 were omitted for clarity, since at this distance the density of bulk compartment has been achieved.

**FIG. 13.**
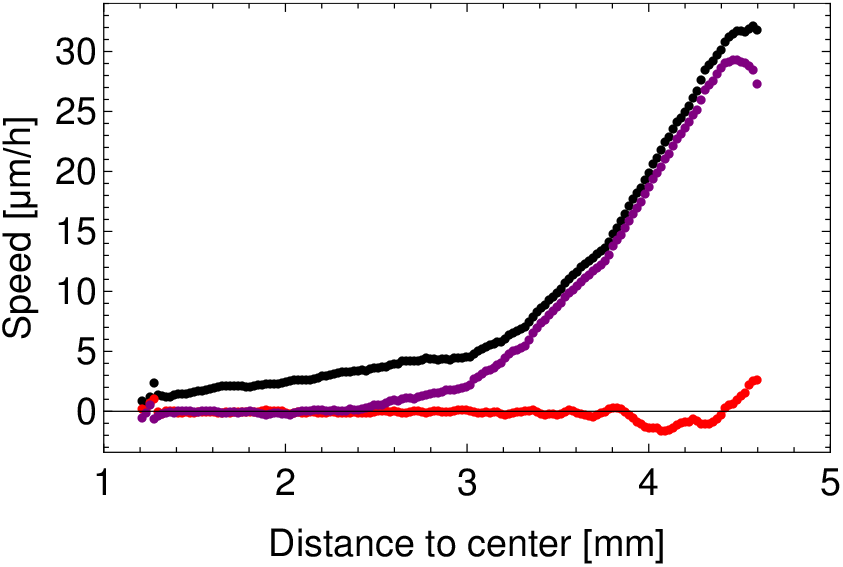
Cell velocity and its components inside the tissue and saturation of the cell speed at the edge. Through PIV analysis (see Appendix D3), the cell speed (in black) and its components (radial in purple, tangential in red) are obtained.

**FIG. 14.**
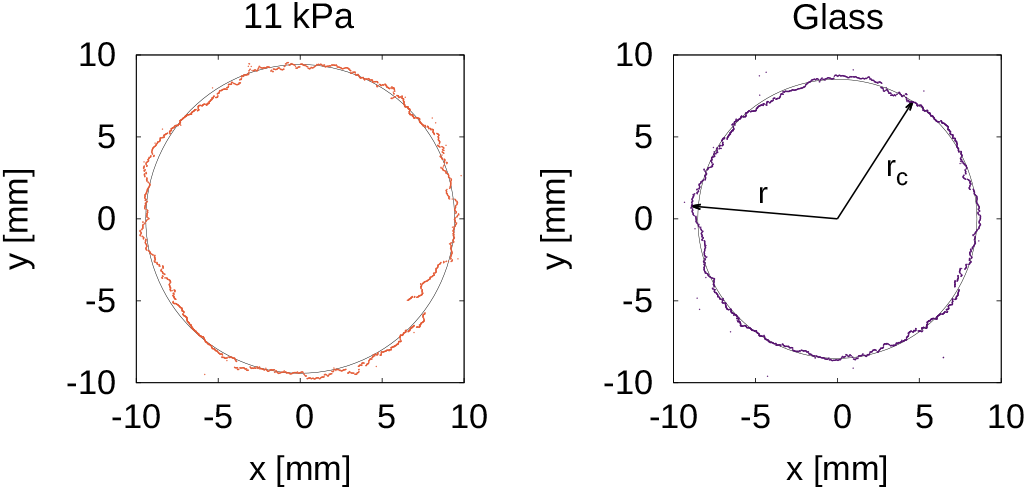
Evaluation of the colony roughness. The edge of the cluster grown on 11 kPa gels and glass respectively and corresponding fitted circle to evaluate the roughness of the border. The vector *r* corresponds to the distance of each pixel to the circle center and *r*_*c*_ is the circle center.

**FIG. 15.**
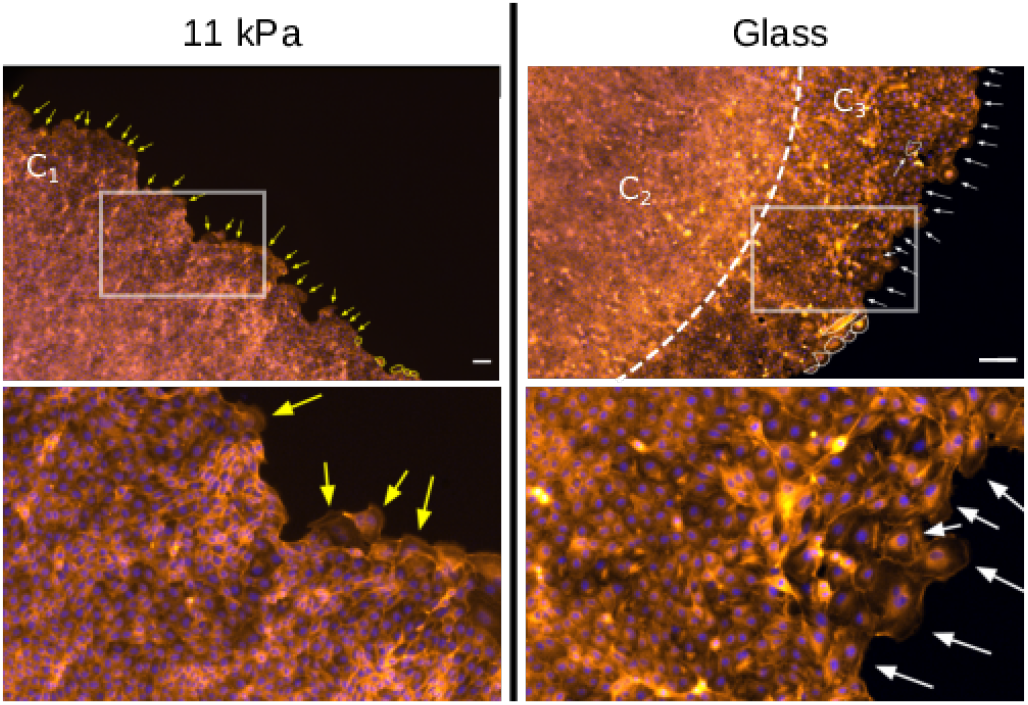
Identifying leading cell phenotype. Leading cells are identified in tissues grown on hard gels (left) and glass (right). The red channel is used for actin, while the blue channel is for the nuclei. Arrows point at leader cells and the scale bars are 100 *μ*m. Leader cells are on average larger on glass substrates (white encirclement on glass (right), yellow encirclement on gels (left)). In contrast to gels, leader cell phenotypes [55] on glass can sometimes be seen deeper inside the tissue and not only on the borderline. The bottom row corresponds to the zoomed-in images corresponding to the highlighted rectangles in the top row.

We capture this compartmentalization using dissipative-particle-dynamics (DPD) simulations [56–58] (technical details in Appendix E, with parameters values summarized in Tab. 2)). The DPD simulations integrates various contributions in the effective terms in a similar way as the theory for the cell shape [14]. As such, the substrate stiffness is accounted for through the friction associated with propulsion over the substrate. The later contribution is a combination of adhesion and traction forces, which are indeed substrate dependent. To obtain the compartmentalisation as seen on glass, the observation of the saturation of the cell velocity at the edge of the colony (see Fig. 13) is incorporated into the model (see Fig. 17). Without this assumption *C*_2_ and *C*_3_ cannot be delineated with the basic model. However, imposing the saturation of cell velocities at the edge in *C*_3_ we find that *C*_2_ spontaneously appears (bottom right panel in Fig. 3b). In turn this can mechanically stabilize *C*_1_ during the spreading of the colony.

**FIG. 16.**
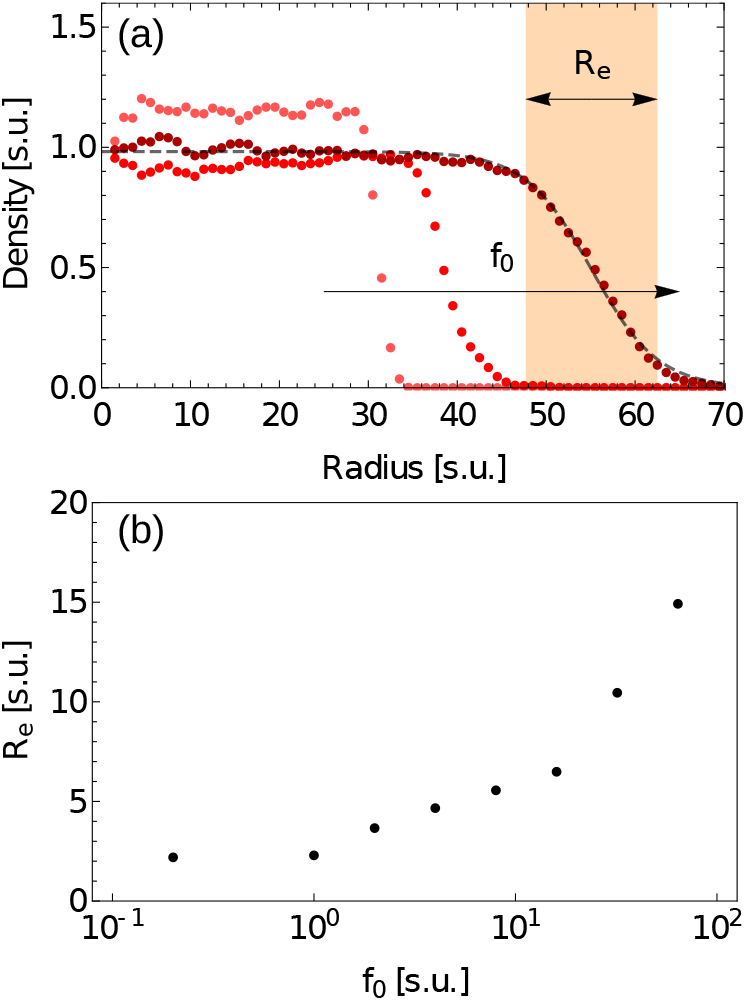
Influence of the parameters *f*_0_ on the shape of the density profile in simulations. (a) Density profiles for different values of *f*_0_ at the same time step. From lightest to darkest: *f*_0_ = 0.2, *f*_0_ = 16 and *f*_0_ = 64. All other parameters and also the initial conditions are identical. The black line on the dark red curve is a fit using a hyperbolic tangent which allow to extract the size of the edge (orange box). Note that contrarily to the results displayed on Fig. 3, no constrain on the velocity of cells at the border has been applied therefore not reproducing the *C*_2_ and *C*_3_ compartments. (b) Size of the edge *R*_*e*_ as a function of the parameter *f*_0_ in semi-logarithmic scale.

**FIG. 17.**
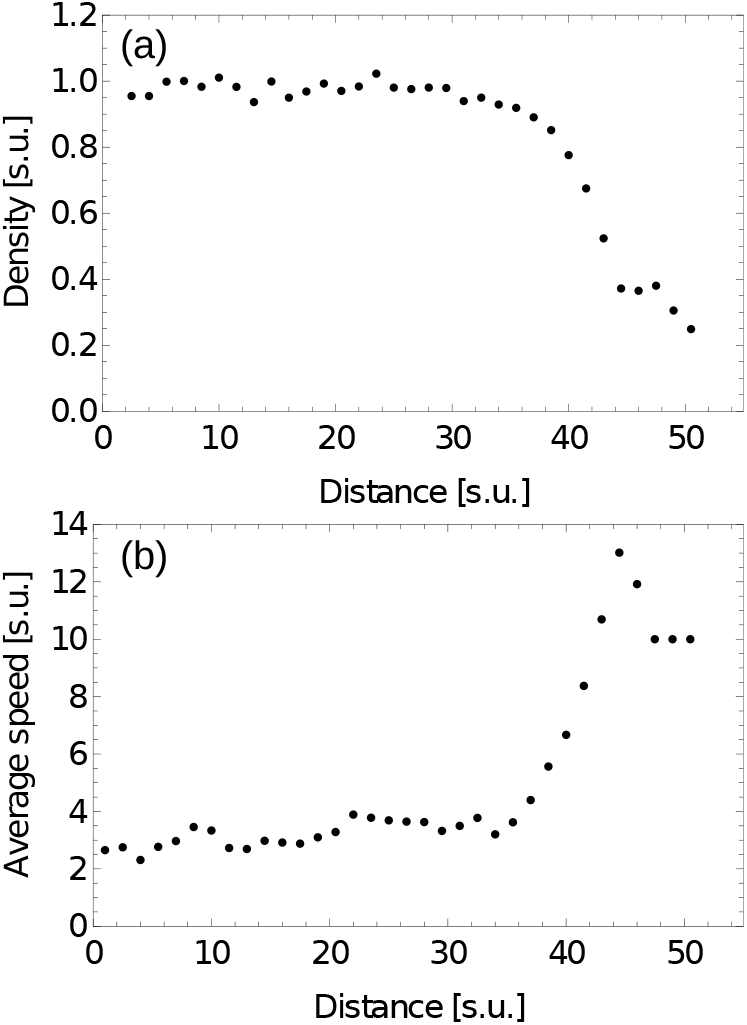
Compartmentalization in the colony as emerging in the simulations, analogous to the glass substrate. In simulations, by externally saturating the cell speed in a ring at the periphery of the tissue, a local modulation of the cell density is seen at the edge. (a) modulation of the density and apparition of the *C*_2_ and *C*_3_ compartments at the tissue edge. (b) Cell speed in the tissue and saturation. The saturation is set to 10 s.u. in a 5 s.u. thick ring.

A different internal structure of the colony has been observed in tissues grown on hard gels. While not displaying a *C*_2_ and *C*_3_ compartments, the colony still possesses an outer compartment similar to *C*_1_ and extending all the way to the edge. On hard gels, only the first outer layer of cells is decorated with the leading cells phenotype (see Fig. 15). Furthermore, we find that large scale roughness (see Appendix D4) is more important on hard gels compared to glass, where roughness emerges mostly from small deviations from the average shape (Fig. 3f). This is consistent with frequent and fast changes of neighbor-hood in the *C*_3_ compartment on the glass (see Fig. 15). On gels, on the contrary, large deviations from the average circular shape have the time to develop since the restructuring of the very edge is slower.

The key new feature in tissues grown on hard gels is the new compartment denoted as H_P_ (Figs. 3a,d,e), in which cell proliferation is inhibited (Fig. 3d,e) but the locomotion is still ongoing (see Fig. 12). Surprisingly, the cell density in H_P_ is significantly higher than in the already discussed, fully arrested H_PL_ homeostatic state (Figs. 1c and 3b). This result is unexpected given the usually reported inverse relation between tissue density and average cell velocity [59]. Note also that the H_P_ compartment can also be seen in the actin signal as shown in Fig. 3e. Furthermore, since H_PL_ and H_P_ display statistically equivalent topologies, this result shows that there could be a more complex relation between tissue structure and motility than previously reported (see Fig. 10).

To further investigate the origin of this highly cooperative state, we use DPD simulations and show that similar behavior appears if a surplus active pressure is generated at the outer edge of H_PL_ (Fig. 3b). We rationalize this surplus active pressure by the higher speed of cells on soft substrates than on stiff ones (see [60] and references therein). Hence, we change the activity of the cells and increase the parameter Γ, which is the amplitude of the force produced by all cells with respect to the direction of the velocity vector, and the value of this parameters is constant across the whole tissue. Its amplitude for the gels is thus set to 1.5 s.u., compared to glass, where it has been chosen as Γ = 0.5 s.u. Indeed, this higher pressure, given its definition, increases the cell speed. This sole difference is sufficient to produce the non-monotonic density profile characteristic for hard gels. The density overshoot in the *H*_*P*_ compartment therefore comes from the competition between activity and friction at the substrate interface. Consequently, the colonies on gels grow somewhat faster than on glass, which indeed seems to be the case even in experiments.

The growth is naturally supported by the proliferation of cells, that by and large dominates the *C*_1_ compartment in gels and *C*_1_ and *C*_2_ on glass (Fig. 3e), but strongly decay toward the very edge of the colony in *C*_3_. This suggest that the probability for division strongly couples to the cell size, yet the proliferation pressure in the colony is built up millimeters away from its edge. The force therefore comes from the core extending the colony. This is different than the force than the force that a non-adhesive interface applies forces inwards and confine the colony [61]. The overall pressure is than spatially dependent and is built from the propensity of cells divide and to move to establish a particular density profile.

In the H_PL_ state proliferation is small, and often occurs at the position of 20-40 *μ*m large defects in the tissue (see Appendix F and Fig. 18). These defects are associated with extrusion events, which start to take place typically 4-5 days after seeding as the homeostatic state is established. In agreement with previous reports in the literature [62], these extrusions do not trigger the appearance of a new colonies. Actually, as the tissue grows beyond the edges of the initial seeded drop, no secondary colonies are observed alongside the main one. Cell divisions thus act to heal the tissue. Given that the extrusions are significantly more common on glass than on gels, residual EdU signal is also more intense. These defects are significantly different to domes [63] which are however significantly larger (average size of the order of 4000 *μ*m^2^). Domes appear as transient blisters when the homeostatic state is achieved (see Fig. 19) and cells start to perform their physiological role, pumping ions from the apical to the basal side of the tissue. Due to the porosity of gels, pumped ions do not accumulate below the basal side of the tissue, and therefore domes do not appear on the gels. However, the extrusions are common and a natural part of the maintenance of the homeostatic state.

**FIG. 18.**
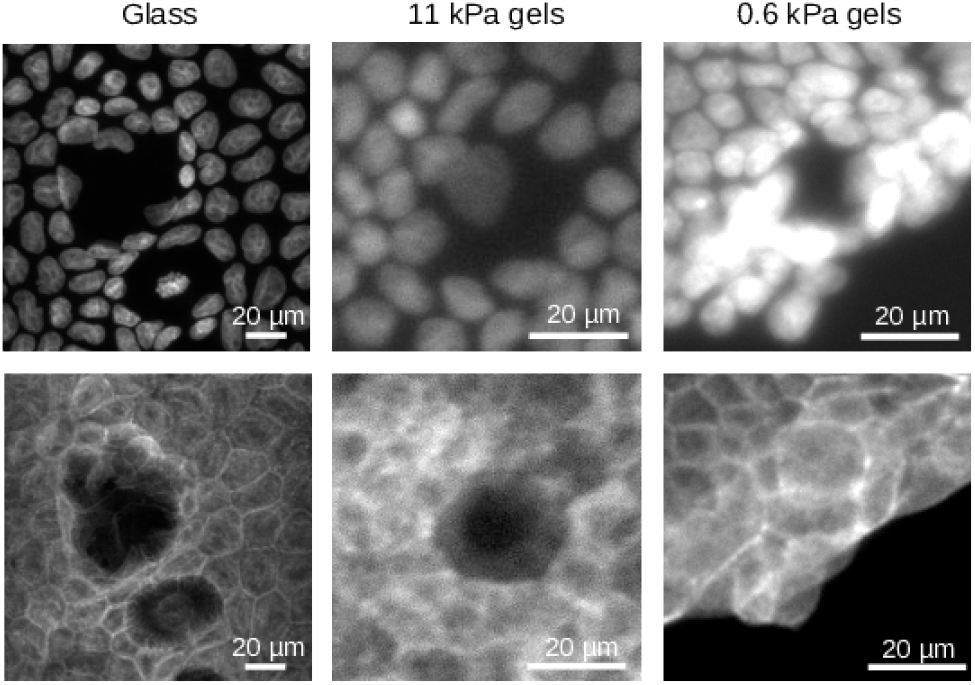
Scars left by extrusion events in the tissue. Pictures of scars left by extrusion seen through nuclei pictures (top) and actin pictures (bottom) for cells grown on glass, hard 11 kPa gels and soft 0.6 kPa gels.

**FIG. 19.**
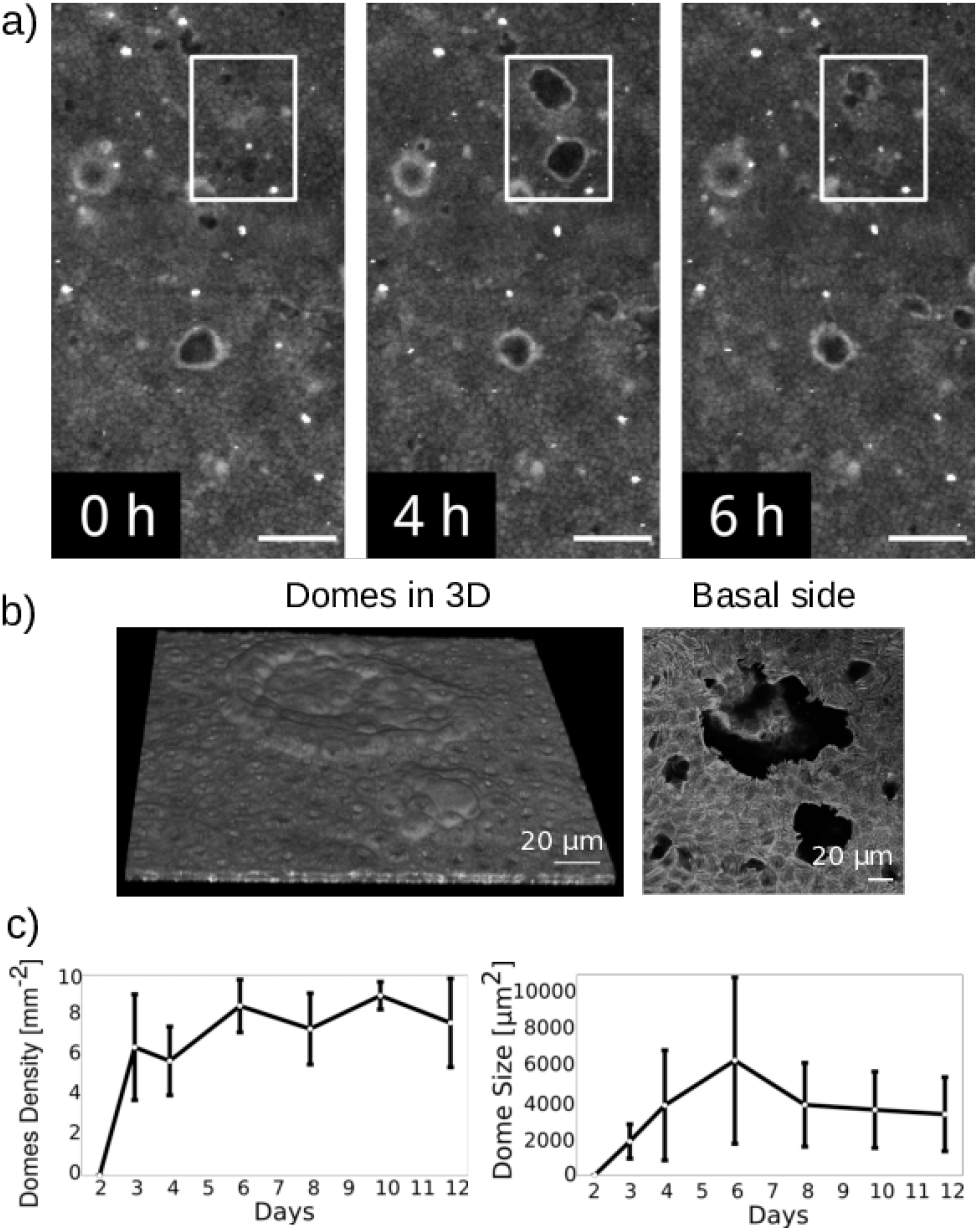
Domes formation within the epithelial tissue. (a) Creation and dissolution of dome-like structures in the bulk of mature MDCK-II monolayers. At 0h, all cells are in contact with the surface. At 4h, the domes are formed in the middle of the field of view. The cells are moved upwards and away from the focal plane. Only the base of the domes is visible in the form of two bright circular structures. At 6h, the circular structures disappear as the domes collapse and the cells return to the focal plane. (b) 3D structure of the domes and corresponding magnified picture of the basal side of the monolayer. (c) Long-term characterization of the domes average density within the monolayer and size.

## V. Macroscopic Organisation on Soft Gels

A fundamentally different mode of compartmentalizing the homeostatic state takes place for soft gels (stiffness of 0.6 kPa). Here, the cells first form a small spherical agglomerate and, over time, create the H_PL_ monolayer in the center of the colony. Consistently with previous work [34], we find a critical minimum size of the microcolonies of 4.7 × 10^−3^ mm^2^ at which the formation the monolayer is observed in the central region, with densities up to 25000 cells/mm^2^ (Fig. 1b). As the monolayer spreads up to 2.8 × 10^−2^ mm^2^, the density decreases until the finite density of about 12580 cell/mm^2^ is achieved in the central compartment of the colony (Fig. 4b). This suggest that finite size effects have a range of about 40 *μ*m, and spread over about 10 cells.

The columnar H_PL_ state (Fig. 1b) is encircled only by a 40 ± 10 *μ*m-thick girdle of cells comprising a few cell layers (top panels in Fig. 4a). Moreover, those girdle cells that are in contact with the substrate build a thick actin filamentous belt that surrounds the whole colony (bottom panels in Fig. 4a). In small colonies, the actin belt is more circular, presumably mechanically stabilizing the central single layered compartment. This compartmentalization into the girdle and the monolayer is maintained over all colonies. For larger clusters, the overall shape is irregular, and the actin belt may vary in thickness. (Fig. 4b).

These highly irregular shapes of colonies emerge from the dynamics of growth and the formation of sub-colonies following cell extrusions. Namely, on soft gels, functional cells are expelled as soon as the monolayer forms, leaving small scars often surrounded by a multilayered ring of cells that is subsequently filled (see Fig. 18). At the same time, small secondary colonies appear and are found throughout the experiment, up to 12 days. Extruded cells land near the mother colony, and develop daughter colonies. As the colonies grow, they merge contributing to the complex shape of the large colony. The merging events are associated with the restructuring of the girdles. This induces line-like defects of smaller monolayer densities (Fig. 4b). Since all colonies present clearly only two compartments, we presume that the reorganization takes place immediately as the two unstructured multilayered compartments start to touch. Given that the locomotion is prohibited everywhere but in the proliferating girdle, this regime is associated with a very different growth dynamics [38], which should be explored in the future.

Finally, a coexistence of the two regime of growth is observed on gels with the stiffness of 1.2 kPa (see Fig. 8). Here, we observe the compartmentalization characteristic for soft and hard gels within a single colony. However, given the uncertainties in the gel stiffness at 0.6 ± 0.3 and 1.2 ± 0.3 kPa, we cannot distinguish if the two coexisting states are result of the fluctuations in the gel stiffness or a genuine coexistence of dynamic phases in a very narrow range gel stiffness. Furthermore, specific consideration should be given to the interplay between mechanosensitivity and adhesiveness. Therefore, it would be interesting to systematically modulate the density of adhesion molecules on the surface, which could be done using existing protocols [21, 64, 65]. Manipulating the density of adhesive sizes was found important in the case of single cells [21, 64], and could influence the position of the transition. Notably, however, it is interesting that the strongest mechano-response of MDCK-II tissues occur exactly in this range. Namely, it is well established that there is a critical range of elasticity at which an individual cells exhibit a mechanosensitive response, often correlated with the stiffness of cells themselves [21]. Individual MDCK-II cells show mechanoresponse in the between 0.6 to 5 kPa, while no significant differences were noted on 11, 20, 34 kPa [34]. That coincides with the range reported for *in vivo* stiffness of kidneys [66, 67] and stiffness of MDCK-II cells themselves [68]. This range also coincides with the range at which we observe major changes in tissue organization and mechanoresponse.

## VI. Discussion

Besides shedding new light on the emergence and the properties of the homeostatic state, the results obtained herein can be used to reconcile some of the apparently contradicting results discussed in the literature. Namely, the mechanoresponse in tissues is truly a cooperative effect involving a significant number of cells. Consequently, clusters containing up to three cells [29] are presumably dominated by the mechanoresponse of individual cells, and not representative of the response of the ensemble. Clusters containing three to seven cells, in which no effect of substrate elasticity was found [29], are potentially too small and the fluctuations are too large to clearly capture changes in cell morphology. Nonetheless, even at these ensemble sizes, the biochemical response clearly couples to the mechanoresponse. For example, treatment of small MDCK-II clusters with TGF-*β*1 growth factor induced apoptosis on soft substrates (<1 kPa), while on hard substrates (>5kPa) it led to the epithelial-mesenchymal transition [33].

Similar effects of cluster size were identified in fibroblasts and in epithelial cells seeded on hard gels and glass substrates. In small aggregates, effects of substrate elasticity were reminiscent of single cell results [30], although under the same conditions, epithelial cell sheets showed no appreciable morphological response [31]. The same behavior has been reported for bovine aortic endothelial cells in confluent layers [32]. Notably, in those conditions, our own samples have shown a large variability. Actually, prior to homeostasis, changes in the distributions of morphological parameters within a single colony are significantly larger than the differences between distributions emerging from different homeostatic states. The exception is the homeostatic state observed on very soft gels, due to the very different growth patterns that are characteristic for growth below the non-equilibrium phase transition [34, 35]. Last but not least, we note that if our colony meets a mechanical or chemically imposed edge, the homeostatic state will expand over the entire surface. The other compartments are thus dynamic, mechanoresponsive structures associated with development of the tissue. Consequently, the same tissues grown in systems of a few hundreds of manometers in size do not show the H_P_ compartment [61, 69]. Nevertheless, cell density can reach values up to 10^4^ cells/mm^2^, showing a mechanoresponsive behaviour, where the H_PL_ state is accurately recovered.

Strong fluctuations prevail even in the homeostatic state, as evidenced by the error bars in Fig. 1c and Fig. 2b. This demonstrates that mechanosensitivity of tissues must be investigated statistically. However, on hard gels (3-30kPa), despite the averaging over homeostatic states occupying tens of square millimeters, we were not able to observe appreciable changes in the colony shape, confirming previous reports [31]. Nonetheless, the manipulation of cohesive and adhesive forces within MDCK-II cell clusters induces changes in the cell spreading areas and proliferation rates, showing that these systems are still sensitive to the cooperative generation of stress [36]. Nevertheless, significant differences between samples grown on hard gels and glass are observed only when a 2mm-thick H_P_ on gels as well as 0.5mm-thick *C*_2_ and *C*_3_ compartments on glass fully form about 4 days after seeding, which again points to the necessity for large sample sizes in studying cooperative behavior of cells in the tissue, and the associated steady states.

## VII. Conclusion

To summarize, we clearly demonstrate that tissue homeostasis depends on the mechanical properties (i.e. Young’s elastic modulus *E*) of the environment on multiple length-scales. The observed self-organization produces several non-equilibrium phases, including a motile, non-proliferating compartment that is of higher density than the homeostatic state. The homeostatic state itself is found to undergo a non-equilibrium phase transition from columnar to squamous tissue with increasing the stiffness of the underlying matrix, as theorized previously [14]. Surprisingly, while the 3D cell shapes and densities change drastically during homeostasis on different matrices, the topology of the steady state is preserved. This suggests that homeostasis can be associated with a set of mechanically universal states.

It remains to be understood, however, what are the developmental advantages of this particular state. It is also not clear which cellular processes and signalling are involved in its maintenance. The intimate relation between the nuclear distributions and the tissue organisation, as seen through the applicability of the set based Voronoi tessellation suggest that a cooperative shape optimisation [70] may be involved. Yet the physical forces and the biological signalling leading to optimisation to this particular distributions are yet to be analyzed, the result of which may be associated with and relevant to tissue development where mechanosensitivity was already found to be important [22, 23]. The mechanical universality of the homeostatic state should be furthermore verified in vivo where mechanical modulation of the environment occurs over long time periods. Tissues should be able to accommodate for these changes by actively maintaining their topology, and not the cell area, as well as proliferation and apoptosis rate, a fact that should be further explored in a more physiological setting.

## Acknowledgements

This work was in part supported by the Deutsche Forschungsgemeinschaft (DFG) through the collaborative research center SFB 755 “Nanoscale Photonic Imaging”, project B8 (FR, CW), and also supported by Deutsche Forschungsgemeinschaft (DFG) - SFB TRR 305-B05 (DD). We furthermore acknowledge German Research Foundation projects RTG 1962 (ASS, SG, DD) and RTG 2415 (ASS, MH), the ERC StG 337 MembranesAct (ASS, SK, JL, LN), intramural funds by the IZKF, project A80 (DD, SG, DV) and by the Emerging Fields Initiative “BigThera”, which was partly supported by the Staedler Foundation (ASS, DD, DV). We thank the Optical Imaging Center Erlangen (OICE) for their support.

## Author Contributions

ASS conceived the study, designed the experiments with FR, the analysis tools with SK, and the simulations with MH. ASS, FR, DD supervised the work. SK, CW, DV, SG performed the experiments. SK, SG, and MH performed structural analysis of the data using tools developed by SK. JL performed the analysis of topological measures using tools developed together with SK. MH performed the simulations using the code developed by LN. DV performed experiments and the analysis associated with velocity distributions. ASS and MH wrote the manuscript with the help of FR and SK. Critical insight was provided by all authors.

## Appendix A Methods

## 1. Cell Culture and gels preparation

MDCK-II cells were obtained from ECACC, UK (#00062107) and cultured in MEM Earle’s medium (#F0325, Biochrom) supplemented with 5% fetal bovine serum (FBS, #F0804, Sigma-Aldrich), 2mM L-glutamine (#G7513, Sigma-Aldrich), and 1% penicillin & streptomycin (#15070-063, Gibco, LifeTechnologies) at 37°C and 5% CO_2_. Cells were passaged every two or three days before reaching 80% confluence.

Elastic polyacrylamide (PA) gels were prepared as described earlier [34]. In brief, appropriate mixtures of acrylamide (40% solution, BioRad) and bis-acrylamide (2% solution, BioRad) were polymerized by addition of 0.1%(v/v) N,N,N,N tetramethylethylenediamine (TEMED) and 1%(v/v) ammonium persulfate (APS) for 60 minutes at RT on plasma cleaned glass cover slips (No.1, 25mm ∅, VWR) that were pre-treated with 3-aminoproyltriethoxysiliane (APTES, Sigma-Aldrich) for 15 minutes and incubated with a 0.5% solution of glutaraldehyde in PBS (Sigma Aldrich) for 30 minutes. For quality control, the Young’s modulus *E* was measured macroscopically by a bulk controlled with a strain rheometer (MCR 501, Anton Paar) using a cone and plate geometry and microscopically by atomic force microscopy (MFP-3D, Asylum Research, Santa Barbara). Typical results of both techniques are given in Fig. 5. We found that the standard deviation of the Young’s modulus *E* is usually below 10%. For quality control reasons, we used the stock solutions not longer than 3 months stored at 4°C and protected form light and ideally prepared as many samples from the same batch as needed.

After polymerization they were washed extensively with PBS and subsequently coated with Collagen-I (BD Biosciences) at 0.02 mg/mL in a 50 mM HEPES buffer using the bi-functional cross-linker Sulfo-SANPAH (Pierce, Thermo Scientific) activated for 10 minutes with UV light (365 nm). Its homogeneity was tested and demonstrated on numerous occasions in [64, 71, 72]. In the current work, we use the conditions optimized in [64] (see Fig. 2c inset therein), at which the concentration of collagen on the surface is fully saturated, to avoid deviations in available collagen density, and to make sure that adhesive properties of the surfaces are identical in all cases.

One series of experiments involved all gels (0.6, 3, 5, 11, 21 kPa) to be prepared concurrently. For quality control of the PA gel, stock solutions stiffness was measured regularly with a bulk rheometer (e.g. see Fig. 5a) as well as by AFM on the ready made PA gels on coverslips (see Fig. 5b). Collagen-I was bought in several bottles from the same batch. All MDCK-II cells came from the same passage and were pooled before seeding to ensure a homogeneous population. All colonies were seeded at once, cultured together under identical conditions, and treated in the same way during the experiment.

## 2. Characterization of cellular features - Fluorescent Staining and Microscopy

We characterize the cells constituting the mechanoresponsive tissues by imaging the following structures and features:
- **Focal adhesions**, which are macromolecular protein complexes located at the basal side of the cell. The main function is the reinforcement of the anchoring of the cells via integrins to extra cellular matrix (ECM) proteins. Besides mechanical linkage it is involved in mechanosensing of the environment and the transduction of signal to the inner of the cell. When binding to ECM proteins (such as collagen, fibronectin, etc.), focal adhesion comprise **Paxillin** in the intracellular compartment. The latter is a small protein (≈65kDa). It acts as an adapter for the recruitment of many other proteins to the cell membrane during adhesion to ECM and mechano-transduction.
- **Actin cytoskeleton**, which is the main scaffold of the cell responsible for contractile stresses. It is built from actin and actin binding proteins (e.g. cross-linkers), which is a protein organized in filaments, bundles and networks. Actin filaments are mostly located at the cell cortex and form together with myosin II mini filaments stress fibres at the basal and apical side of the cells, the “cellular muscles”, that exhibit contractile forces. Ventral stress fibers (at the basal side of the cells) are connected on both ends to focal adhesions.
- **Dividing cells** using 5-ethynyl-2-deoxyuridine (EdU), which is a thymidine analogue that can be incorporated into DNA during its replication. EdU is an indicator that cells underwent the S-phase of the cell cycle.

For imaging, tissues were fixed using a 10% solution of formaldehyde (formaldehyde, #47608, Sigma-Aldrich) in PBS for 5 minutes, and the cells were permeabilised for 10 minutes using a 0.5% solution of Triton X 100 (Carl Roth). After washing with PBS, samples were blocked using 3% BSA (#A9418, Sigma-Aldrich) in PBS for 30 minutes at RT, which preceded another Triton X 100 treatment of 5 minutes at RT, followed by 3 washing steps with PBS.

Filamentous actin was stained using Phalloidin Atto 550 (#AD550-81, Atto Tec) by incubating with a solution [1:250] in 3% BSA in PBS for 1.5h at RT. *β*-catenin was stained by incubating with the primary antibody (mouse, #C7082, Sigma-Aldrich) solution [1:200] in 3% BSA in PBS, for 2h at RT. This was followed by the secondary anti-mouse IgG Alexa Fluor 488 (goat, #SAB4600388, Sigma-Aldrich) [1:250] in 3% BSA in PBS for 30 minutes at RT. Last, nuclei were stained using Hoechst 33342 (H3570, Invitrogen/Thermo Fisher Scientific) [1:1000] in 3% BSA in PBS for 30 minutes at RT. EdU staining was performed using the Click-it Plus EdU Alexa-Fluor 488 Imaging Kit (#C10637, Molecular Probes, Thermo Fisher Scientific) according to manufacturer’s instructions. Cells were incubated with 10*μ*M EdU for 4h. Paxillin staining was performed with a primary antibody anti-Paxillin (rabbit, #SAB4300384, Sigma-Aldrich) [1:1000] in 3% BSA in PBS for 2h at RT followed by the secondary antibody anti-rabbit IgG Alexa Fluor 488 (goat, #AP132JA4, Sigma-Aldrich) [1:250] in 3% BSA in PBS for 30 minutes at RT. Samples were mounted using Fluoroshield histology mounting medium (#F6182, Sigma-Aldrich, USA) on cover glasses (26 × 76mm, “Menzel Glaeser”, #1, Thermo Fisher Scientific).

Epifluorescence microscopy images were acquired on an inverted microscope (Zeiss Cell Obsever Z1) using 5× and N-Achroplan 20× objectives, using AxioCam M3 and AxioVision software package (all Zeiss). Confocal microscopy was performed on a Leica LSM SP5 laser-scanning microscope equipped with a white light laser and a 63× and a 100× oil immersion objective yielding fields of view of (246 *μ*m × 245 *μ*m) and 100× (155 *μ*m × 155 *μ*m) respectively. The step size in the z-direction was kept constant at 0.25 *μ*m.

All tissues in grown in one series were fixed and stained simultaneously and imaged consecutively without changing the microscope settings.

## Appendix B Homeostatic state - Imaging

## 1. Contact inhibition of proliferation and locomotion in homeostatic states

To demonstrate the contact inhibition of proliferation, we search for cell divisions using EdU staining of H_PL_ state on glass, hard and soft gels (see Fig. 6(a-c)) following the above described protocol. To demonstrate the inhibition of locomotion, PIV analysis (vide infra) was performed and shows only slow positional fluctuations of cells (see Fig. 6(d-f)).

## 2. Density in the homeostatic state on glass and hard gels

On glass and hard gels, we first evaluate the cell density *ρ*_0_ in the central region of the clusters (2.3 mm radius around cluster center, tiled into 267*μ*m × 575.6*μ*m fields of view). Standard deviation *σ*_*p*_ is evaluated as the deviation in cell density obtained from different fields of view. Hence, *σ*_*p*_ is a measure of fluctuation densities in the sample. The homeostatic state is defined as all fields of view showing a density in the range of *ρ*_0_ ± 2*σ*_*p*_. The H_P_ state state on gels is determined as all fields of view in which the density is higher than *ρ*_0_ ± 2*σ*_*p*_ (see Fig. 7). The average cell area is calculated from the mean cell density. The latter was obtained from hand corrected segmented images of cell nuclei (accuracy above 99%), and estimated using a MATLAB routine discussed in details in our previous publication [34].

**TABLE 3.**
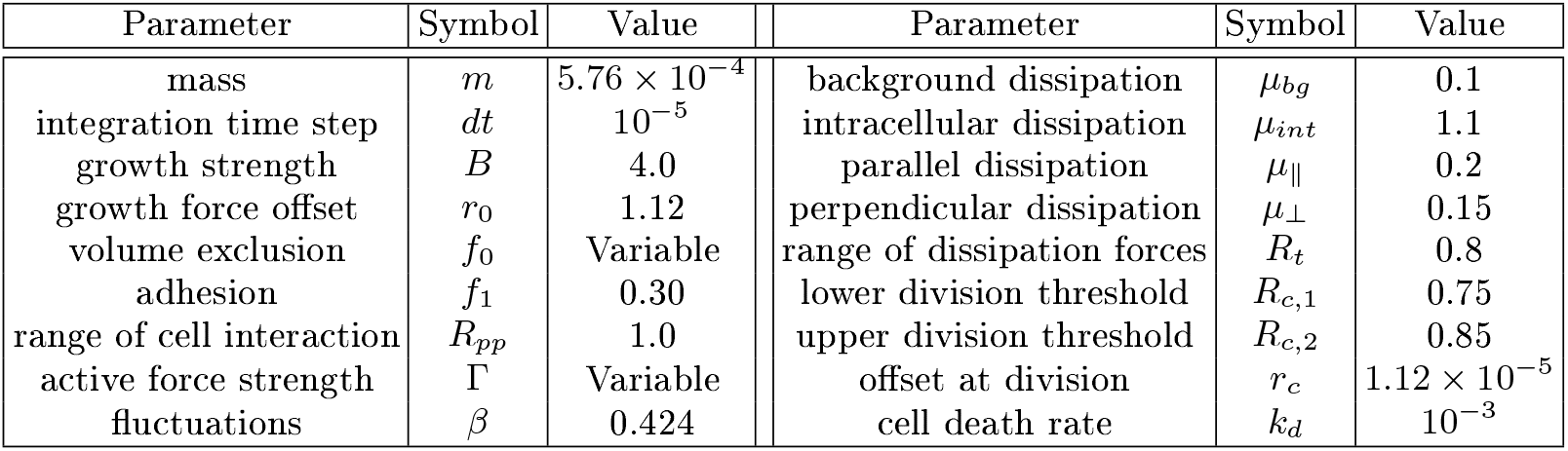
Parameters used in simulation and expressed in simulations units (s.u.). Parameters listed as *variable* see their values change through the article.

**TABLE 2.** Quality of SVT-extracted measures. Relative average deviation of the of the SVT-based morphological measures from the measures obtained via *β*-catenin stained, segmented and hand-corrected pictures of the membrane, of fully tubular cells within the H_PL_. As discussed in our previous work these deviations are comparable or smaller than deviations that are inherent to uncorrected *β*-catenin segmented images [38].

| Substrates | Area | Perimeter | Elongation | Neighborhood |
| --- | --- | --- | --- | --- |
| Soft Gels | 6.62% | 4.20% | 6.90% | 3.77% |
| Hard Gels | 6.86% | 5.10% | 8.97% | 6.64% |
| Glass | 10.54% | 5.52% | 10.27% | 6.92% |

**TABLE 1.** Information about the amount of cells considered in obtaining the density of the homeostatic states.

| Stiffness | $\rho(H_{PL})$ (cells/mm<sup>2</sup>) | Number of cells | $\rho(H_P)$ (cells/mm<sup>2</sup>) | Number of cells |
| --- | --- | --- | --- | --- |
| 0.6 kPa | $12.58 \times 10^3$ | $500 \times 10^3$ | | |
| 3 kPa | $7.71 \times 10^3$ | $26 \times 10^3$ | $8.31 \times 10^3$ | $23 \times 10^3$ |
| 5 kPa | $7.12 \times 10^3$ | $34 \times 10^3$ | $7.56 \times 10^3$ | $5 \times 10^3$ |
| 5 kPa | $7.08 \times 10^3$ | $45 \times 10^3$ | $8.49 \times 10^3$ | $31 \times 10^3$ |
| 11 kPa | $7.37 \times 10^3$ | $56 \times 10^3$ | $8.81 \times 10^3$ | $26 \times 10^3$ |
| 21 kPa | $7.14 \times 10^3$ | $57 \times 10^3$ | $8.00 \times 10^3$ | $14 \times 10^3$ |
| glass | $6.45 \times 10^3$ | $10 \times 10^3$ | | |
| glass | $7.11 \times 10^3$ | $9 \times 10^3$ | | |
| glass | $7.02 \times 10^3$ | $8 \times 10^3$ | | |

With regards to the density in the range *E* 〉 [3 : 21] kPa, the average densities for the homeostatic states *ρ*(H_PL_) and *ρ*(H_P_) (when relevant) are reported in Tab. 1.

When reporting the density in the H_P_ state in Fig. 1, the averages and the standard deviations ate found from the entries in Tab. 1. Notable, this variance is very similar to *σ*_*p*_: it is somewhat larger on glass than on hard gels, and it is the largest on soft gels. With these data (comprising more than 850 000 cells) we cannot establish statistically different results for average densities of the *ρ*(H_PL_) and the *ρ*(H_P_) on hard gels.

## 3. Density of colonies grown on soft gels

Following the previously established criteria (see [34]), colonies smaller than 2.8 × 10^−2^ mm^2^, are classified as small. Below this size, we presume that finite size effects still play a role in the monolayered structure. For the analysis of the density in small clusters, we use the entire monolayer. In the analysis of the large clusters density, averaging was performed over a significant sections of the monolayer for which there is no other compartment in the same field of view. The same criteria were applied in determining the topology related measures, which were extracted only for large clusters (size > 2.8 × 10^−2^ mm^2^). The scars left by extrusions (see Appendix F) are removed manually from the images since cells cannot be counted in these areas. Here, scars left from the merging of colonies has very little effect, if any.

## 4. Cell volume measurements

Samples of cell monolayers grown on 0.6 kPa, 5 kPa and glass substrates were stained for actin and imaged with confocal microscopy (see details Appendix A). The cell height was estimated 4 days after seeding to make sure there is a small amount of defects wihtin the monolayers. The density of the tissue segments are calculated by counting cells. The following values have been obtained: On glass 6546 cells/mm^2^, on 5 kPa gels 6944 cells/mm^2^, in the large cluster on 0.6 kPa gels 14760 cells/mm^2^ and in the small cluster on 0.6 kPa gels 23010 cells/mm^2^. For hard gels and glass, the densities are somewhat smaller than the average reported for the H_PL_ state, yet within two standard deviations of density fluctuations *σ*_*p*_ used to define the state. In this way we are able to balance the accuracy of two independent measurements - density and height.

The cell height is given by the mean intensity of the actin signal in the stacks. These stacks are plotted as a function of the z-axis position as shown Fig. 1. The corresponding images have been taken at day 4 after seeding instead of day 6 to avoid any defect within the tissue that typically arise when the tissue ages. This also leads to slightly smaller densities that the one reported in Fig. 1(c), as it is indicated in the labels in Fig. 1(d). More information about the time evolution of the density in the homeostatic state can be found in Fig. 12. The samples showed characteristic two maxima of actin signal corresponding to the positions above and under the cell nuclei (see Fig. 1d). Intensity at the beginning (*I*_*bottom*_) and the end (*I*_*up*_) of the cell monolayer is estimated via the two inflection points of the curves. To this method, we associate to the z-positions an error of ±5% of the intensity at the inflection point divided by the gradient at the inflection point. This method therefore associates larger z-axis estimation errors to the slowly decaying actin intensity profiles. The cell area is estimated from the density at in the homeostatic state on the corresponding gels.

The average cell volume is obtained as a product of the mean cell area and the mean cell height. The deviations of the volume are calculated by propagating the respective errors in the cell area and height.

## Appendix C Homeostatic state - Characterization

## 1. Morphological characterization

To examine the properties of the cells and their nuclei, we select in each tissue a number of regions of interest from the original 20×-magnified images (with a pixel size of 0.31*μ*m), in order to avoid the effects of defects and domes on glass. All regions were collected in the homeostatic state, far enough from the external compartments. On glass we sampled 35 regions of interest (areas between 14000-78000*μ*m^2^) providing 2575 cells for morphological analysis. On 5 kPa in total 9095 cells were analyzed from 24 regions (28000-128000 *μ*m^2^). On 11 kPa PA gels we have 5244 cells from 27 regions (28000 and 128000 *μ*m^2^). All morphological measures were obtained using a previously described procedure [34, 40], from the Set-based Voronoi Tessellation (SVT). Due to the balance of forces, this tessellation is particularly suitable for analyzing the homeostatic states, as the error in determining all morphological measures is between 5% and 8%, depending on the cell size in comparison to pictures of *β*-catenin stained tissues. A more in-depth description of the precision of this method for the densities discussed in this paper can be found in Tab. 2. Pictures illustrating the quality of the tessellation with respect to the *β*-catenin stained pictures can be found in Fig. 9.

Error bars in Fig. 2b have been obtained by considering the distributions of 50 statistically independent subsets of cells. For *n* cells considered on a given substrate, statistically independent subsets of *n*/2 cells were considered. The distributions of cell area and perimeters, nucleus area and perimeters, and nuclei and cells elongation have been computed for those 50 subsets always using the same binning. For each bin, the average value and standard deviation of those 50 iterations were computed. The error bars displayed in Fig. 2b are centered around the average value and the size of the bars are given by ± the standard deviation of all 50 average values.

## 2. Topological characterization

To describe the topological properties of tissues grown on substrates of different stiffnesses, we focus on three different geometric laws. The first law, the Aboav-Weaire’s law, linearly correlates the number of neighbors of cell with *n* sides and the average number of neighbors *m*(*n*) that cells adjacent to ones with *n* sides have [41–43]. Mathematically, it writes

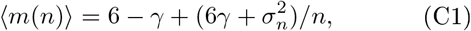

where 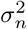 is the second central moment of the distribution of *n*, and *γ* is a constant. The two other laws describe the linear relationship between the number of neighbours *n* and the average area 〈*A*(*n*)〉 of a cell with *n* neighbours:

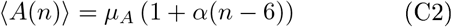

Here, *μ*_*A*_ is the average area of the cell in the assembly and is a constant. A similar expression, namely,

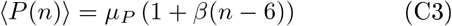

exists for the average perimeter 〈*P*(*n*)〉 of cells with *n* neighbours and is known as Desh’s law [53].

These law are illustrated in Fig. 10, for the H_PL_ tissues on all substrates, and also for the H_P_ state observed on hard 11 kPa gels.

## Appendix D Analysis of the colony structure

## 1. Density profiles from high resolution imaging

On glass and hard gels, after seeding a droplet, a confluent monolayer is formed from 12 hours after seeding. Consequently, small groups of cells may exist only in the very early stages of the experiment, and are a likely consequence of the initial seeding. As the colony grows beyond the edges of the initial seeded drop, no secondary colonies are observed alongside the main one.

Cell density profile throughout cell colony is analyzed after fixing the samples 4 or 6 days after seeding. High precision measurements are obtained from images of cell nuclei, imaged at magnification of 20×. All images were segmented and hand corrected to achieve fidelity of 99%, as described in detail in previous work [38].

In fully reconstructed clusters, imaged at 5 magnification, density profiles are extracted from the calibrated fluorescence intensities. The grids of images were stitched using the Fiji^3^ plugin based on the work published by Preibisch *et al* [74]. Cell density distributions were estimated from the stitched images of the Hoechst stained cell nuclei. First, the mean intensity was determined in 51*μ*m wide circular rings drawn around the geometric center of the cluster (see Fig. 11). Consequently, linear mapping of the mean intensity to the mean cell density is performed for each tissue individually, after benchmarking based on at least 20 different segments with an area of at least 0.5mm^2^, where the cell number and fluorescence intensity could be evaluated with very high accuracy.

## 2. Dynamical analysis of the homeostatic states

Locomotion within the H_P_ state is demonstrated by imaging its position relative to the center of the tissue on day 4 and day 6 as shown in Fig. 12.

## 3. Particle Image Velocimetry (PIV) analysis and speed profiles

To determine velocity profiles, tissues were grown for 6 days on glass. For this picture 10,000 MDCK-II cells were seeded in a dense droplet into a collagen-I coated well of an IBIDI 2-well glass chamber slide. 6 days after seeding, cell imaging was performed on a Zeiss AXIO Observer.Z1 microscope with a moving stage and the incubation chamber capable of long term cell incubation. For fluorescent visualization, the cluster was stained with CellTrace Violet live cell dye (Invitrogen). Time steps were 15 minutes apart and the duration of the complete time lapse was 6h. In total, 25 time points were sampled (24 time steps between them). Full image was stitched out of 9 fields of view (3 × 3 and 5% overlap in X & Y direction). Fields of view had individual sizes 1.4 mm × 1.4 mm and the pixel size of 2.76 *μ*m.

After imaging, a PIV analysis was performed with an ImageJ plugin as described in [75]. PIV interrogation windows were squares 44 *μ*m (16 px) in size, correlated with search windows in the subsequent time step of 88 *μ*m (32 px) size. Gird distance between interrogation windows was 22 *μ*m (8 px, 50% overlap).

Visualization of the PIV calculated velocity field was achieved in post-processing. Within a separate subsection of the ImageJ PIV plugin, a vector field plot was created by using the originally calculated velocity text file as input. The vectors plotted had their starting point positioned at the (x, y) coordinate. Vector direction was the direction determined by the (*V*_*x*_, *V*_*y*_) velocity components. Arrow color and arrow length were depicted depending on the speed magnitude.

For the analysis of the homeostatic state (Fig. 6), PIV analysis has been performed on a region of interest of a square of 4 mm × 4 mm, subdivided in PIV interrogation windows of 44 *μ*m^2^. The latter were correlated via search windows of 88 *μ*m^2^. The gird distance between interrogation windows is of 22 *μ*m therefore leading to an overlap of 50%. A total of 25 images taken at 15 minutes intervals have been considered, giving 24 velocity matrices. The average of those 24 matrices gives the results given in Fig. 6.

## 4. Roughness of the colony edge

To calculate the distribution of roughness, first the edge is determined using an home-made algorithm applied on the thresholded images. The algorithm finds the first non-zero intensity pixel on lines drawn from the center of the images to the pixels of the border of the images. This gives the coordinates of the pixels of the edge of the tissue. On the ensemble of coordinates of each pixel describing the edge, we fit a circle with variable center and radius, which well describes the average colony shape, as shown in Fig. 14. Consequently, the distance of each edge pixel to the center of the circle *r* is determined, and the deviation from the circle radius *r*_*c*_ recorded in the histograms displayed in Fig. 3.

## Appendix E Simulations

The numerical model is adapted from previous works [56–58]. Each cell is made of two points which are sources and targets of all forces in the system. The “nucleus” of each cell is taken as the geometrical center of these two points. Finally, cell shape and geometrical features are obtained by Voronoi Tessellation based on the position of all nuclei.

The forces applied to building particles are a growth force 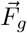 in order to trigger cell division, a cell-cell inter-action 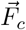, an active motile force 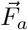 and a random force for inherent fluctuations 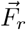. The expression for these forces are the following

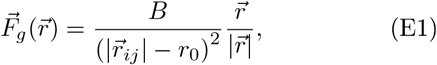

where the force only applies between building particle within a single cell only, *B* and *r*_0_ are parameters and 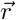 is the vector pointing from one building point to the other,

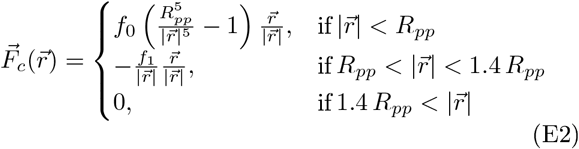

where the force applies between building particles of different cells only, *f*_0_, *f*_1_ and *R*_*pp*_ are parameters and 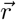 is the vector pointing from one building point to the other,

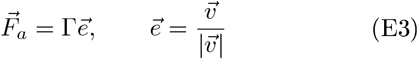

where Γ is a parameter and 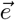 is the unit vector pointing in the direction of the cell nucleus speed, defined as the average speed of its two building particles,

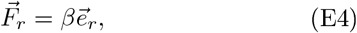

where *β* is drawn from a Normal distribution with zero mean and standard deviation *σ* while 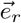 is a random unit vector with an orientation drawn from an uniform distribution. These two last forces are applied to all building points in the system.

Dissipation within the simulations are accounted for thanks to the following forces. For the dissipation with the background, each building particle is submitted to

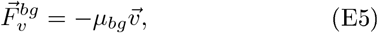

where 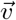 its velocity. Dissipation within a single cell is accounted for by the force

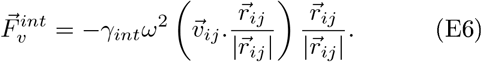

Dissipation in the relative motion of cells is decomposed into a direction parallel and perpendicular to the line joining the building particles of each cell involved. One has

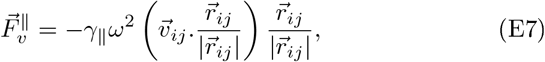

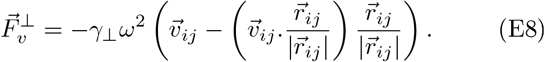

The weight function *ω* appearing in the previous formula

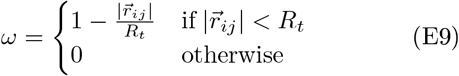

The equation of motion for each building particle are

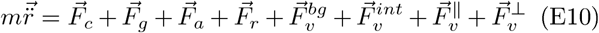

These equations are integrated for each cell using an Euler algorithm with time step *dt*.

Cell division as well as cell death is included in the simulations. Cell death consists in the removal of a random cell at rate *k*_*d*_ *dt* at each time step. Cell division is based on the distance between the two building particle 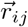 for all cells. At each time step, all cells have the following division probability

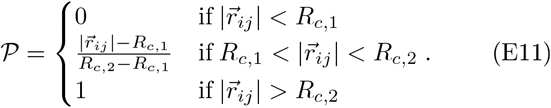

When division occurs, the two daughter cells inherit one building particle of their mother. Their second building particle is created at a distance *r*_*c*_ with random uniform orientation with respect to the first building particle. The velocity of the daughter cells building particle is 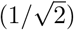 times the velocity of their mother counterparts.

The shape of cell is based on the Voronoi tessellation of the plane, using the particle nuclei position as generating point. Given the position 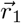 and 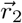 of the building points, the nuclei position is defined as 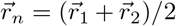. The algorithm is hand-made. This technique allows for the extraction of the cell area and cell perimeter (for associated formula, see [38]).

Finally, we give in Tab. 2 the value of the parameters used in the simulations with the exception of Γ (see 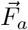) and *f*_0_ (see 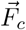) which are varied in this article. We also provide in Fig. 16 an analysis of the influence of the parameter *f*_0_ on the shape of the density profile, and in Fig. 17 a comparison between the speed profile and density profiles obtained from simulations when a constrain on the speed is applied. We choose here a speed saturation to 10 s.u. in a 5 s.u. thick ring at the edge of the tissue.

## Appendix F Defects in the monolayers

As the tissues mature, different defects can be found in the monolayer, which are intrinsic features of the homeostatic state. Present on all substrates and the first to appear are scars associated with extrusion events (Fig. 18), in which cells are ejected from the tissue. The characteristic size of these defects is 20-40 *μ*m. The tissue repairs these scars, often by divisions of neighbouring cells.

About 8 days and after seeding, cells deposit a significant amount of their own extracellular matrix. This induces local inhomogneities in adhesiveness, and the appearance of different types of defects on the monolayer in the central (oldest) compartment of the colony. Furthermore, we observe a drop of average density and significant inhomogeneity in cell and nuclei structures.

On glass, a prominent type of a defect are domes, which emerge due to the physiological role of the MDCK-II cells to pump from the apical to the basal side. On non-porous substrate, this induces blisters in the tissues due to the accumulation of ions on below the basal side [63]. Domes appear in large quantities after the homeostatic state is achieved. However, these are not permanent structures and after some time, they collapse back to the surface and the comprising cells adhere again (see Fig. 19(a,b)). The size of the domes, their life time and frequency of occurrence (Fig. 19c) is related to the the content of the medium. In the conditions used in the current experiments, the characteristic size is 4000 *μ*m^2^.

